# Pocket-based molecule generation with an SE(3)-equivariant language model leads to a potent and selective HPK1 inhibitor with *in vivo* efficacy

**DOI:** 10.1101/2025.09.23.678079

**Authors:** Bin Xi, Han Wang, Guanglong Sun, Bowen Zhang, Ruihan Mao, Yuyang Ge, Yang Wang, Jiangtao Zhang, Yiting Pan, Feng Zhou, Yuji Wang, Zhenming Liu, Daohua Jiang, Huting Wang, Wenbiao Zhou, Bo Huang

**Author notes:** These authors contributed equally to this work (Bin Xi, Han Wang, Guanglong Sun, Bowen Zhang).

## Abstract

Deep learning shows promise in structure-based drug discovery, yet challenges persist in generating pharmacologically plausible molecules with valid 3D conformation and decent binding mode in the pocket. We introduce SE3-BiLingoMol, an SE(3)-equivariant Transformer for pocket-based 3D molecule generation, addressing two key limitations of existing language-model approaches. First, it uses Geometric Algebra Transformers for SE(3)-equivariant handling of continuous 3D coordinates. Second, a bidirectional attention mechanism mitigates conformational errors accumulated during autoregressive sampling. These innovations enable SE3-BiLingoMol to generate 2D drug-like, 3D geometrically valid molecules with superior binding modes. Validated on DUD-E dataset containing over 100 targets, the model achieved state-of-the-art performance in *de novo* design and optimization. We applied SE3-BiLingoMol to design potent and selective inhibitors for HPK1, a promising immunotherapy target. Through an iterative human-AI workflow, integrating AI generation with experimental validations (X-ray crystallography, bioassays), we identified Cmpd. 6. This novel tetracyclic compound demonstrates potent HPK1 inhibition, excellent cellular activity, favorable pharmacokinetics, and robust anti-tumor *in vivo* efficacy as monotherapy and with PD-1 blockade. Our work establishes a sophisticated generative AI framework for 3D molecule design and demonstrates its application in developing cancer immunotherapy.

**Highlights:**

● **SE(3)-Equivariant Language Model for Molecule Generation:** We introduce a dual-channel SE(3)-equivariant language model that supports *de novo* molecule design and optimization, addressing key challenges in prior language model-based 3D molecule generation approaches.
● **Discovery of Potent HPK1 Inhibitors:** Application of this model led to the discovery of highly selective HPK1 inhibitors with robust *in vivo* efficacy, resulting in a promising lead compound.
● **AI-Guided Rational Drug Development Paradigm:** This work establishes a paradigm for AI-guided rational drug development, showcasing AI as a “co-pilot” that inspires drug designers with novel ideas in an “evidence-based context”.

## Introduction

Structure-Based Drug Design (SBDD)^1,2^ is a well-accepted paradigm in drug discovery that leverages the three-dimensional (3D) structures of biological targets to facilitate the rational and efficient design of novel molecular entities within a defined chemical space. This approach primarily aims to generate molecules satisfying three key criteria^3^: (1) high binding affinity to specific target sites, (2) novel molecular scaffolds exhibiting desirable drug-like properties (indicative of efficient chemical space exploration), and (3) well-conformed 3D poses with minimal internal strain. Traditional *in silico* methods, such as virtual screening^4,5^, explore molecules within predefined chemical spaces^6,7^. While these methods can ensure reasonable pocket binding modes and valid 3D conformations through the application of well-established force fields and stereochemical empirical rules, they often encounter challenges in enhancing the efficiency of chemical space exploration, thereby restricting the novelty achievable within predefined chemical spaces^8^.

The application of deep generative models has revolutionized the efficiency of chemical space exploration^9,10^. For instance, several studies^11–15^ have successfully demonstrated the rapid generation of novel molecules with drug-like topologies using representations such as 1D SMILES/SELFIES strings or 2D molecular graphs^16–18^. Despite these advancements, such early approaches frequently lack explicit consideration of biomolecular structures in 3D space. Consequently, their capacity to generate novel molecules with optimal interactions with binding pockets is inherently limited^19^, thereby precluding their complete fulfillment of all three aforementioned SBDD criteria.

To address this critical challenge, 3D molecule generative models have garnered increasing attention within the SBDD community. These models typically comprise two principal components: a 3D feature aggregation module and a 3D molecule generative module. Thus, recent developments in this field can be analyzed from two perspectives: methods for feature aggregation and strategies for molecule generation. Feature aggregation methods primarily fall into two categories: graph-based and language model-based approaches. Similarly, molecule generation strategies are broadly classified as autoregressive-based or diffusion-based methods.

Graph-based methods are more commonly employed for feature aggregation compared to language model-based approaches, largely due to the existence of well-established architectures capable of acquiring SE(3)-equivariance. SE(3)-equivariance is crucial because models lacking this property treat the same 3D structures with different spatial rotations and/or translations (i.e., symmetry transformations) as distinct samples during training. This deficiency hinders the effective learning of intrinsic molecular spatial properties, such as bond lengths, bond angles, or torsion angles, which should be insensitive to the symmetry transformations of the entire molecule. This ultimately leads to the generation of problematic 3D conformations and suboptimal interactions with the binding pocket during inference^20,21^. Advances in geometric deep learning^22^ have provided mature solutions for graph-based models to achieve SE(3)-equivariance. These graph-based feature aggregation methods can subsequently be integrated with either autoregressive or diffusion strategies for 3D molecule generation. For example, some studies^23–25^ have proposed autoregressive prediction of atomic positions, atom types, and chemical bonds, while others^26,27^ have utilized diffusion strategies to generate molecules by iteratively denoising a sample drawn from a prior distribution. However, despite their effectiveness in capturing 3D spatial relationships among atoms and generating molecules with valid 3D conformations that exhibit decent binding modes, these graph neural network (GNN)-based models have been reported to produce molecules with flawed 2D topologies and limited drug-likeness^28^. Examples include molecules containing a high proportion of *sp*^2^ carbons, honeycomb-like flat ring arrays, or an inappropriate number of rings (either too many or none)^28^.

Conversely, language models, while excelling at generating molecules with desirable 2D drug-likeness, often struggle with fulfilling criteria related to 3D conformational validity and achieving proper pocket binding. This limitation stems from their less mature support for handling continuous 3D coordinates in an SE(3)-equivariant manner. Specifically, these approaches leverage Transformer architectures^29^, originally designed for natural language processing (NLP), to extract chemical and structural features from sequential representations of molecules and proteins. While the Transformer’s success in learning “grammatical” rules in NLP has fostered expectations for its ability to improve the topological validity and drug-likeness of generated molecules, the intricate and continuous nature of atomic coordinates poses significant challenges for modeling SE(3)-equivariant 3D protein-ligand complexes. Although some attempts^30,31^ have been made to discretize Cartesian coordinates into discrete tokens, these methods frequently lack accurate capture of critical binding geometries and remain sensitive to symmetry transformations. For instance, Feng et al. (2022)^30^ proposed a fragment-based SMILES representation that incorporates ring information to enhance the learning of molecular topologies using a language model, while treating 3D conformation prediction as a discrete multiclass classification problem. Their model reportedly improved the proportion of drug-like molecules generated with valid 3D conformations through autoregressive sampling. Nevertheless, they still face challenges in handling continuous atomic coordinates in an SE(3)-equivariant fashion. Additionally, the frequently employed autoregressive sampling in language models can suffer from issues such as error accumulation, leading to poor-quality molecular conformations and limited molecular diversity. For example, some literature^27^ reports that such methods are prone to mode collapse, resulting in the generation of molecules with inaccurate distributions of bond lengths for various single and double bonds compared to reference sets.

Considering that current graph-based models, regardless of their generation strategy (autoregressive or diffusion), frequently struggle to consistently improve drug-likeness, and language models often lack SE(3)-equivariance while suffering from error accumulation associated with their typically coupled autoregressive generation strategies, there is an urgent need for novel methods. Such methods must ensure both the drug-like topology of generated molecules and an enhanced perception of spatial structures. Logically, this could be achieved by either refining graph-based feature aggregation techniques or by introducing SE(3)-equivariance and anti-error accumulation mechanisms into language models.

To address these challenges, we present SE3-BiLingoMol, a novel SE(3)-equivariant language model designed for structure-based molecule generation in 3D space. SE3-BiLingoMol employs an encoder-decoder framework, where the encoder processes protein pocket structures as inputs while the decoder autoregressively generates 3D molecules binding to the pocket. It incorporates a dual-channel perception mechanism that separately perceiving atomic coordinates (vector features) and scalar attributes (e.g., atom types, bond types), and allows them to interact with each other by leveraging a geometric algebra Transformer^32^ as an E(3)-equivariant backbone architecture. Instead of relying on rule-based constraints or post-fixing adjustment on the conformations, SE3-BiLingoMol allows free exploration of conformational space, and enable a self-refinement of ligand conformations through a bidirectional attention mechanism to alleviate the 3D conformational errors accumulated during generation process. This tactic improves the conformation validity by around 18%. SE3-BiLingoMol was trained on a curated protein-ligand complex dataset that containing 53,871 ligand-protein complex structures. Its performance was evaluated on 102 targets from the Directory of Useful Decoys-Enhanced (DUD-E) dataset^33^, benchmarking against state-of-the-art (SOTA) methods. The results showed that SE3-BiLingoMol not only outperformed these models across various metrics, but also exhibited inference speed that is more than one order of magnitude faster than others, highlighting its substantial potential for advancing drug design.

We demonstrated SE3-BiLingoMol’s practical impact through its pivotal role in the discovery and optimization of potent and selective inhibitors of Hematopoietic progenitor kinase 1 (HPK1). HPK1 is a critical negative regulator of T-cell receptor signaling, making it a promising yet challenging target for cancer immunotherapy due to the need for high potency and selectivity^34,35^. We initiated the discovery campaign with an existing compound which displayed moderate activity but sub-optimal selectivity and toxicity. Employing an iterative human-AI workflow, SE3-BiLingoMol was instrumental in guiding both the generation of novel core scaffolds, and the subsequent optimization through targeted fragment generation. This rational design process, continuously refined by experimental feedback from crystallography and *in vitro* assays, led to the successful identification and optimization of a novel tetracyclic series, culminating in the discovery of Cmpd.6. This highly promising candidate exhibits strong potency and selectivity, improved safety profiles, and good *in vivo* efficacy in tumor models, validating the transformative potential of SE3-BiLingoMol within a synergistic AI-driven drug discovery paradigm.

## 1. Results

### 1.1. SE3-BiLingoMol Model Development

#### 1.1.1. Model Construction

SE3-BiLingoMol is an SE(3)-equivariant, structure-based model designed for autoregressive generation of molecules with 3D conformation using protein pocket structures as input. The architecture and sampling process of SE3-BiLingoMol is illustrated in Figure 1A-B (also see Methods Sec. 3.1-3.4 for details). Our model integrates a pocket encoder and a ligand decoder by leveraging Geometric Algebra Transformers (GATr) layers^32^ as the core building blocks. It supports separate encoding of scalar features (e.g., atom types, bond types etc.) and vector features (i.e., spatial coordinates) while maintaining SE(3)-equivariance.

**Figure 1.**
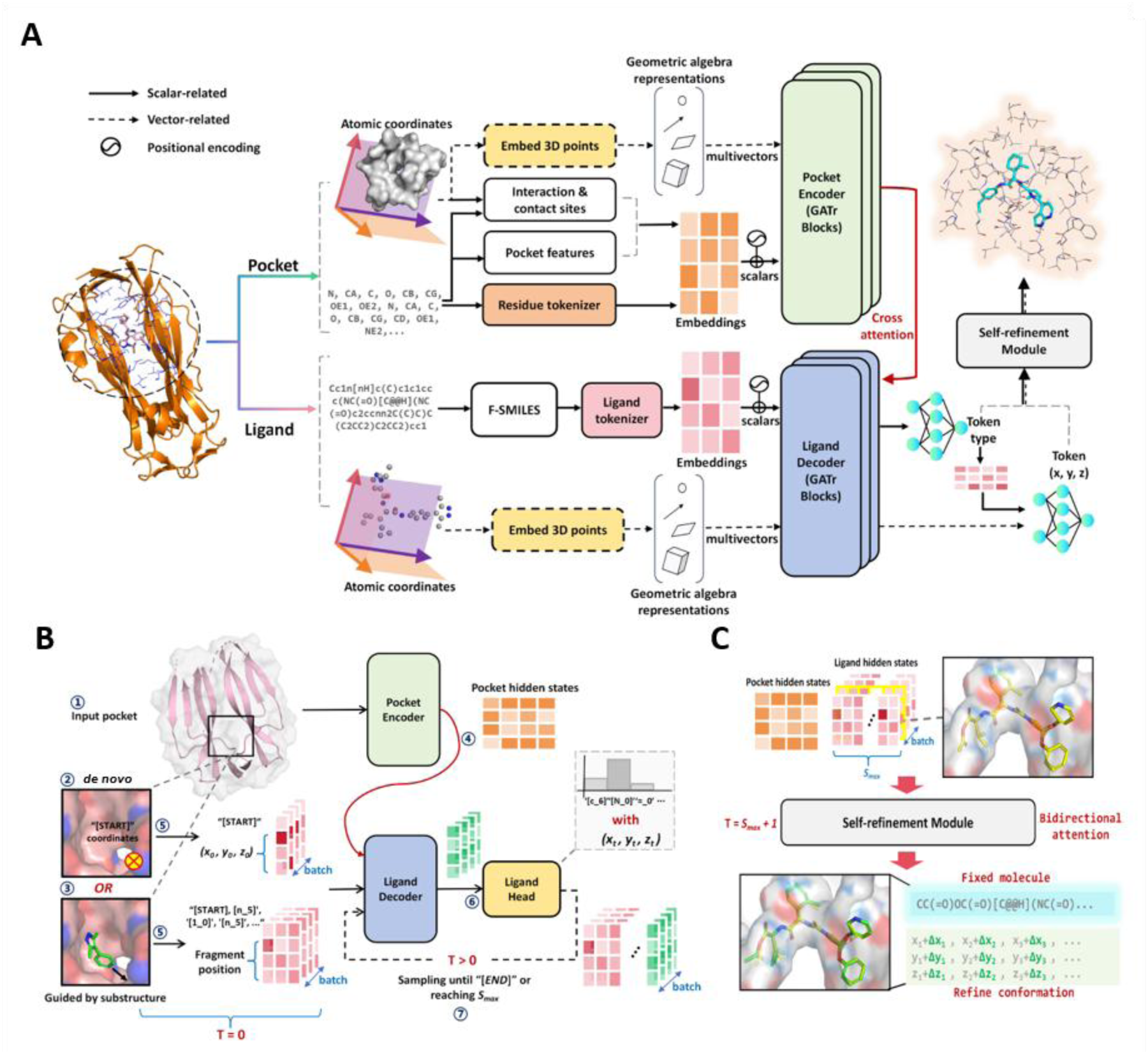
Overview of the SE3-BiLingoMol model. **(A) SE3-BiLingoMol architecture**. SE3-BiLingoMol comprises a pocket encoder, a ligand decoder and a self-refinement module, with GATr E(3)-equivariant layers as key building blocks. It separately encodes scalar features (e.g., atom types, bond types), and vector features (i.e., spatial coordinates) represented as multivectors using projective geometric algebra. Solid and dashed arrows show the flow of scalar and vector feature information, respectively. **(B) Autoregressive sampling strategy of SE3-BiLingoMol.** This panel depicts the cyclical workflow for generating molecules, applicable to both *de novo* compound discovery and substructure-guided molecule optimization. **(C)** T**he self-refinement module.** This module architecture is simply a copy of ligand decoder but using a bidirectional attention mechanism, which enables each token to be aware of other tokens (both ligand and pocket). The self-refinement module is only used after a molecule is sampled, and it is designed to fix the molecule topology (and atomic types) and only allow the atomic positions to be relaxed.

Prior to encoder input, pocket information is prepared into four distinct representations. These include vector-related residue atom positions defining spatial coordinates, and scalar-related residue atom tokens, pocket features, and interaction and contact site indicators. Pocket features encompass hydrogen-bond donor/acceptor indicator, hydrophobicity, aromaticity, formal charges, and residue types. Interaction and contact sites are labels indicating residue atoms prone to forming non-covalent interactions (NCIs) with potential ligands (Details provided in Methods Sec. 3.3 and Supplementary Materials).

The ligand decoder models a conditioned distribution of ligand tokens with continuous atomic positions from the encoder’s pocket context. Specifically, a ligand molecule is encoded into molecule tokens by using fragment-SMILES (FSMILES) representation^30^, which utilizes a ring-first traversal approach to reorganize SMILES sequences into ring units with ring types and connection sites labeled explicitly. Feng et. al.^30^ demonstrated that such representation enhances the perception of molecular geometry especially for language-based generative models.

Spatial coordinates for pocket and ligand atoms are embedded as 16-dimension multivectors, and then passed into separate GATr blocks, where a multivector is a representation of a 3D vector by the projective geometric (or Clifford) algebra (see Methods Sec. 3.1-3.2 or refreneces^32^ for details).

In addition, a learnable self-refinement module was designed by leveraging a bidirectional attention mechanism to allow local relaxation of generated molecular conformations in a rule-free manner with molecular topology preserved as illustrated in Figure 1C. This design can largely alleviate conformational errors accumulated in an autoregressive sampling (details provided in Discussion), and thereby ensures robust and geometrically consistent molecular generation.

SE3-BiLingoMol was trained on a curated dataset containing 53,871 protein-ligand complexes from the Protein Data Bank (PDB)^36^. This dataset resulted from a systematic screening of PDB complexes, employing a rigorous protocol based on the PDBbind methodology^37^ (details provided in Method Sec. 3.5). We employed targets from DUD-E for model evaluation. To prevent information leakage during training, a clustering-based filtering approach was implemented. Specifically, the 102 protein targets from the DUD-E evaluation database were combined with the protein structures from our curated training dataset. This combined pool was then clustered using Foldseek^38^, with a TM-score cutoff of 0.5 to define clusters. All collected protein structures that clustered with any of the DUD-E targets were subsequently removed from our final dataset, resulting in the exclusion of 10,224 proteins.

SE3-BiLingoMol is designed for two key applications in drug design: *de novo* compound discovery and substructure-guided molecule optimization. For *de novo* design, the model can generate molecules from scratch using a specified binding protein, making it well-suited for discovering innovative molecular scaffolds. Alternatively, for substructure-guided optimization, the model starts from a provided molecular fragment and grows the molecule, where the provided molecule fragment efficiently narrows the chemical space and enables precise, targeted compound optimization. The detailed description of the sampling strategy supporting these scenarios can be found in Methods Sec. 3.4 and Supplementary Materials Part 1.

#### 1.1.2. Model Evaluation

For comparison with our SE3-BiLingoMol, six structure-based generative models reporting SOTA performance were selected as baselines: Pocket2Mol^24^, TargetDiff^26^, Lingo3DMol^30^, PokcetFlow^39^, PMDM^40^, and MolCraft^27^. Among them, Lingo3DMol is a representative of symmetry non-equivariant language model. Other models are graph-based equivariant models equipped with autoregressive (Pocket2Mol, PocketFlow) and diffusion-based (TragetDiff, PMDM, MolCraft) sampling strategy. For these models, the official implementations were used directly from their GitHub repositories.

The DUD-E dataset was employed for model evaluation due to two primary considerations. First, this dataset comprises 102 distinct protein targets, representing a broad spectrum of biological categories including Kinases, Proteases, G protein-coupled receptors (GPCRs), and ion channels, thereby ensuring good target diversity for robust evaluation. Second, a critical feature of DUD-E is the inclusion of over 200 experimentally validated active ligands with measured affinities for each target. This feature facilitates direct comparison between AI-generated molecules and experimentally validated active compounds, thereby mitigating the limitations associated with relying solely on purely computational metrics for AI generative model assessment.

Therefore, our model evaluation strategy comprised two principal components: computational metrics-based and semi-experimental evidence-based assessments. The computational assessment primarily focused on appraising the 2D and 3D molecular quality of AI-generated compounds, alongside the inference speed of the AI model. Specifically, 2D quality was assessed by evaluating drug-likeness and chemical diversity, while 3D quality was assessed by checking AI-generated 3D conformation validity and ligand-pocket binding modes. In contrast, the semi-experimental evidence-based assessment investigated the model’s capability to reproduce known active compounds. The comprehensive definitions of these metrics are provided in Supplementary Materials Part 2.

Each model under investigation generated approximately 1,000 molecules per DUD-E target. We first assessed the drug-likeness of these compounds. Following Feng et. al.^30^ and Fan et. al.^28^, we defined a drug-like region in chemical space as having a Quantitative Estimate of Drug-likeness (**QED**)^41^ over 0.3 and a synthetic accessibility score (**SAS**)^42^ less than 5. As shown in Table 1, language-based models, including our SE3-BiLingoMol and Lingo3DMol, exhibited a high percentage of drug-like molecules (% **Druglike Molecules**) with values over 80%. In contrast, graph-based models such as Pocket2Mol, TargetDiff, and PDMD showed a lower percentage, with values close to or below 60%. Although also graph-based, PocketFlow and MolCraft exhibited a high percentage of drug-like molecules. PocketFlow achieved good drug-likeness by refining its output through a multi-stage, rule-based post-processing pipeline that applies explicit bond order, valence, and ring topology corrections. MolCraft’s high drug-likeness is partially attributed to its propensity to generate molecules with relatively small molecular weights. Furthermore, when assessed using more specific metrics like non-aromatic ring percentage and maximum fused ring number, MolCraft exhibited drug-likeness issues similar to other graph-based models (details provided in Discussion).

**Table 1.** Summary of evaluation results for molecules generated on DUD-E targets (*n*=102). Metrics with an upward arrow (↑) / downward arrow (↓) indicate higher / lower is better, while these without an arrow are listed for reference. The top two results for each metric are highlighted in **bold** and <u>underlined</u>, separately. Outlying results, defined as µ-1.5σ for metrics where higher is better (↑), µ+1.5σ for metrics where lower is better (↓), and µ±1.5σ for others (where µ and σ represent the mean and standard deviation for the current metric), are highlighted in red. Metrics denoted with an asterisk (*) are computed using a weighted average, with weights corresponding to the number of drug-like molecules for each target.

|  | Pocket2Mol | TargetDiff | Lingo3DMol | PocketFlow | PMDM | MolCraft | SE3-BiLingoMol (Ours) |
| --- | --- | --- | --- | --- | --- | --- | --- |
| # of Generated Molecules | 99250 | 93546 | 100385 | 102000 | 92987 | 101704 | 99911 |
| Mean MW | 443 | 366 | 369 | 311 | 342 | 299 | 373 |
| Mean QED (↑) | 0.46 | 0.50 | 0.53 | 0.52 | 0.47 | 0.62 | 0.54 |
| Mean SAS (↓) | 4.0 | 4.9 | 3.3 | 2.9 | 4.6 | 3.6 | 3.7 |
| % Druglike Molecules (↑) (QED>0.3 & SAS<5) | 60.5% | 48.6% | 82.5% | 80.3% | 51.7% | 86.0% | 83.4% |
| % Aromatic rings | 41.8% | 13.5% | 45.6% | 45.5% | 16.8% | 34.0% | 35.7% |
| % Non-aromatic rings | 32.1% | 48.8% | 11.1% | 12.4% | 43.5% | 37.2% | 15.0% |
| The comparison below involves only drug-like molecules (QED>0.3 & SAS<5) |  |  |  |  |  |  |  |
| # of Druglike Molecules | 60089 | 45425 | 82771 | 81860 | 48108 | 87428 | 83306 |
| Mean MW | 386 | 299 | 347 | 268 | 305 | 285 | 361 |
| Mean QED (↑) | 0.56 | 0.60 | 0.59 | 0.60 | 0.59 | 0.65 | 0.59 |
| Mean SAS (↓) | 3.5 | 4.1 | 3.2 | 2.7 | 3.9 | 3.3 | 3.5 |
| Diversity* (↑) | 0.86 | 0.89 | 0.84 | 0.88 | 0.89 | 0.88 | 0.85 |
| % Conformation Validity (↑) | 61.8% | 41.4% | 60.6% | 43.8% | 65.5% | 86.2% | 87.1% |
| Strain Energy<br>25%, 50%, 75% (kcal/mol, ↓) | 69, 126, 230 | 139, 313, 643 | 13, 40, 177 | 54, 132, 352 | 58, 110, 209 | 20, 47, 102 | 24, 52, 134 |
| Mean Pocket Occupancy* (↑) | 0.70 | 0.70 | 0.58 | 0.57 | 0.50 | 0.54 | 0.76 |
| Mean Min-in-place Score* (↓) | -6.8 | -6.3 | -6.6 | -5.4 | -5.6 | -6.1 | -6.9 |
| Mean Redocking Score* (↓) | -7.6 | -7.1 | -7.7 | -6.4 | -6.5 | -6.9 | -7.6 |
| % Min-in-place < Redocking (↑) | 17.9% | 15.2% | 12.0% | 15.5% | 17.8% | 20.1% | 27.2% |
| % IMP (↑) | 11.2% | 9.6% | 9.2% | 5.9% | 4.9% | 8.5% | 11.7% |
| PLIF Recovery Rate* (↑) | 21.5% | 21.4% | 29.0% | 9.9% | 9.3% | 16.9% | 35.3% |
| % ECFP_TS>0.5 (↑) | 5.9% | 2.0% | 31.4% | 12.7% | 2.0% | 16.7% | 31.4% |

**Note:** This table summarizes the performance of SE3-BiLingoMol against baseline models across key evaluation metrics. These metrics are broadly categorized into two types: **computational-based assessments**, which include 2D and 3D molecular quality evaluations, and **semi-experimental evidence-based assessments**, specifically the Active Reproduction Capability test.

Prior to further evaluations, it is necessary to eliminate molecules of poor drug-likeness, because some previous studies^30^ have demonstrated that including such molecules can compromise pocket binding mode assessment. Specifically, molecules with low drug-likeness can sometimes exhibit deceptively good scores when using force field-based scoring functions for ligand-pocket binding quality assessment. This occurs because such molecules may possess synthetically challenging scaffolds or unfavorable physicochemical properties that, despite their poor drug-likeness, still exhibit favorable shape complementarity and strong polar interactions with the pocket. Therefore, in subsequent analyses, we conducted thorough assessments by only considering drug-like molecules (QED > 0.3 and SAS < 5). For completeness, we also provide the distributions of generated molecules projected onto the entire QED-SAS space in Supplementary Materials Part 3.

In addition to drug-likeness, our computational assessment of 2D molecule quality also included diversity. As shown in Table 1, the AI models under investigation produced drug-like molecules with comparable diversity.

Having assessed 2D molecular properties, we next turned our attention to the three-dimensional quality of the generated drug-like molecules. We first evaluated local conformation quality using the recently reported HEAD method^28^. This method employs machine learning force fields to detect local abnormal conformations based on energetically implausible local environments, achieving near quantum mechanical (QM) accuracy. Our SE3-BiLingoMol outperformed other models in this test. Beyond local conformation, we then evaluated global conformation quality using PoseCheck^43^ to calculate the global strain energy of each molecule. SE3-BiLingoMol also achieved the top ranking in this evaluation. In addition to assessing intrinsic ligand conformations, evaluating ligand-pocket binding quality is also critical. We conducted this assessment across several key metrics (Table 1): the pocket occupancy (**Mean Pocket Occupancy**), the favorability of force-field-based binding scores (indicated by **Mean Min-in-place Score** in Table 1), the superiority of generated binding modes over redocked poses (**% Min-in-place < Re-docking Score**), comparative binding quality against co-crystallized molecules (**% IMP**), and the ability to capture observed non-covalent interactions (NCIs) (**PLIF Recovery Rate**). Across all these evaluated metrics, SE3-BiLingoMol demonstrably outperformed other models, indicating its superiority in handling 3D poses.

Beyond these computational assessments, we conducted a semi-experimental evidence-based assessment by evaluating generative models’ ability to reproduce or generate molecules structurally similar to known active compounds from the DUD-E dataset. Specifically, for each model, we measured the percentage of targets where at least one generated molecule exhibited a Tanimoto similarity (based on ECFP4 fingerprints) greater than 0.5 with any known active compound for that target. This metric is designated as **% ECFP_TS > 0.5** in Table 1. Our SE3-BiLingoMol and Lingo3DMol both achieved a **% ECFP_TS > 0.5** of 31.4%, notably outperforming all other baseline models in reproducing known active compounds.

Finally, we also evaluated the inference speed of each model. Our SE3-BiLingoMol demonstrated exceptional efficiency, generating approximately 500 valid molecules in just 90 seconds (Extended Data Table 1). This represents an acceleration of one to two orders of magnitude compared to other leading models. The high inference speed of SE3-BiLingoMol is attributed to key techniques such as Key-Value cache (KV cache), multi-query attention, and efficient GPU memory utilization, which are detailed in Supplementary Materials Part 4. This notable speed advantage is critical for high-throughput applications in drug discovery.

In summary, SE3-BiLingoMol achieves SOTA performance across multiple metrics while simultaneously offering enhanced generation speed.

### 1.2. SE3-BiLingoMol Application in HPK1 Inhibitor Design

#### 1.2.1. Novel Scaffold Discovery

In order to thoroughly demonstrate the practical application of SE3-BiLingoMol in drug design, we systematically elaborated its pivotal role in the effective design of HPK1 inhibitors. The overall workflow is illustrated in Figure 2. As depicted in Figure 2A, our starting point was an initial hit compound, SW898 reported in patent CN116063329A. As detailed in Supplementary Materials Part 5, SW898 exhibits moderate enzymatic and cellular activity against HPK1. Its selectivity is suboptimal, showing >85% off-target inhibition against a panel of kinases, including GLK, MAP4K5, TBK1, and TNIK. Additionally, SW898 demonstrated certain cytotoxicity in the HepG2 CTG assay. Both this off-target activity and the observed cytotoxicity manifested as a narrow two-fold dose window for IL-2 induction in human primary T cells, as shown in Supplementary Materials Part 5. This two-fold dose window quantitatively describes the dose-dependent profile of small molecule-stimulated IL-2 secretion by T cells. Specifically, for compounds exhibiting a bell-shaped dose-response curve for IL-2 secretion, the two-fold dose window is defined as the concentration difference between the ascending and descending arms of the curve where IL-2 secretion reaches two-fold above baseline levels. A narrow two-fold dose window typically indicates either high compound toxicity leading to cell death or poor target selectivity.

**Figure 2.**
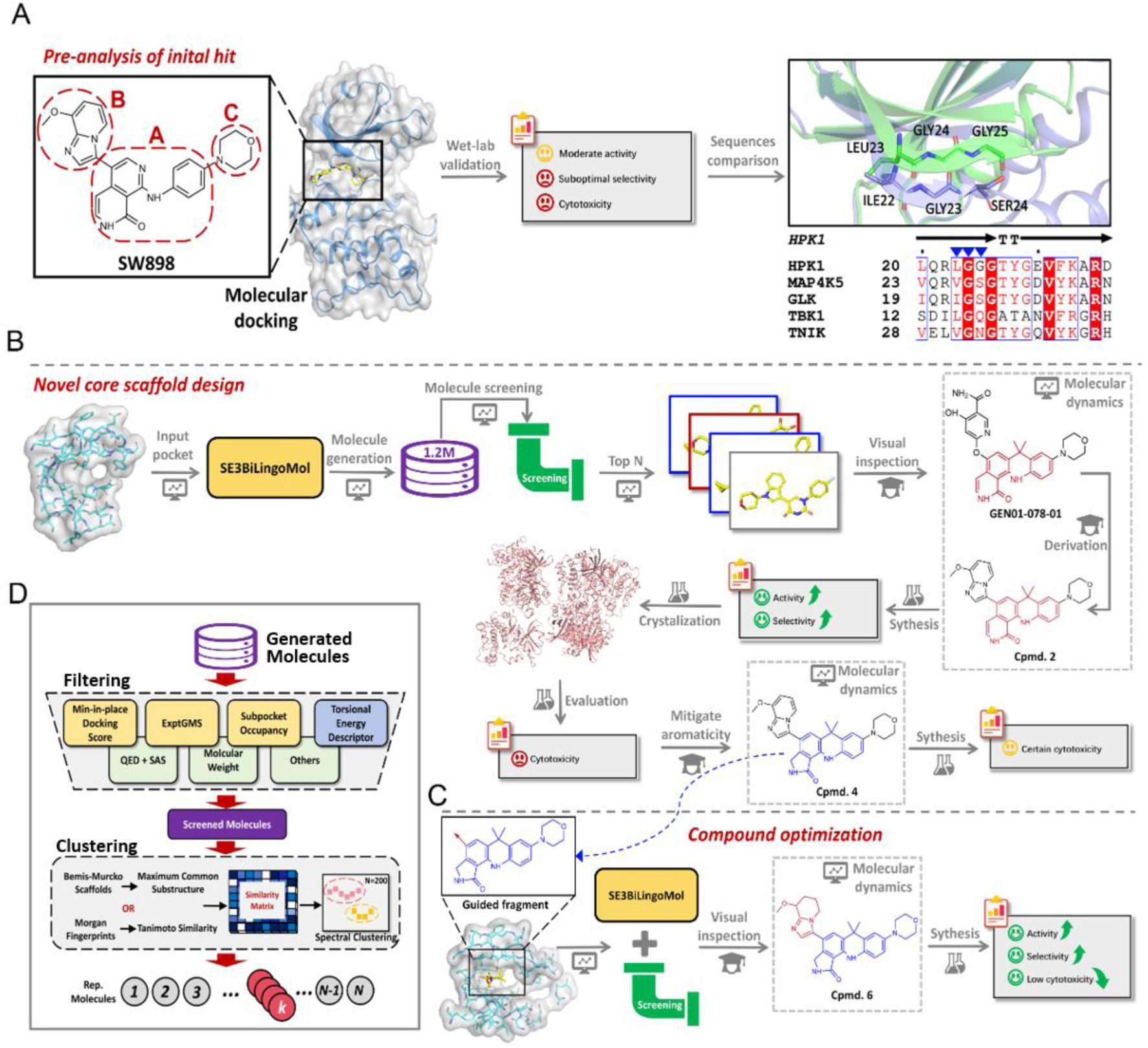
Overall Workflow for SE3-BiLingoMol-Guided HPK1 Inhibitor Design. **(A) Initial hit compound analysis and structural rationale for design.** The starting hit compound SW898 which is reported in patent CN116063329A is divided into three regions (i.e. A, B, and C) for the purpose of clear structural description in subsequent studies. HPK1 and off-target kinases (GLK, MAP4K5, TBK1, and TNIK) displayed sequence variations within the L23-G24-G25 P-loop motif (indicated by blue triangles). **(B-D) Schematic diagram of SE3-BiLingoMol-driven design workflow.** Schematic representation of the iterative design process employing SE3-BiLingoMol. **(B) Novel core scaffold discovery.** This phase starts from an empty pocket and focuses on identifying new molecular scaffolds from scratch. **(C) Compound optimization.** This phase starts with the pocket structure plus a provided molecular fragment and focuses on growing the molecule in the pocket. **(D) Generation-screening paradigm.** Both core scaffold discovery and compound optimization operate within this paradigm, where SE3-BiLingoMol-generated molecules undergo a screening process comprising a filtering module and a clustering module.

Our objective was to leverage the structural information of HPK1 and other kinases to enhance SW898’s HPK1 activity and selectivity while reducing its toxicity. We initiated this by performing a structural analysis. Specifically, we conducted a comparative analysis of the primary sequences and 3D structures of HPK1 and the identified off-target kinases as shown in Figure 2A. This analysis revealed sequence variations within the L23-G24-G25 motif in the HPK1 P-loop across these kinases. For instance, L23 in HPK1 is substituted by isoleucine in GLK or valine in TNIK and MAP4K5. Considering the differences in size and rigidity between isoleucine and valine compared to leucine, we hypothesized that designing molecules with an optimized fit to the L23-G24-G25 region of HPK1 would enhance HPK1 activity and thereby improve selectivity over other kinases.

To facilitate the design of HPK1 inhibitors which can form interactions with L23-G24-G25 region, we employed SE3-BiLingoMol for molecule generation. The workflow comprises novel core scaffold discovery (Figure 2B) and compound optimization (Figure 2C). Both parts followed a generation-screening paradigm, where SE3-BiLingoMol supports molecule generation, and various filters and clustering modules facilitate the subsequent screening (Figure 2D).

The workflow for novel core scaffold discovery is shown in Figure 3A. The binding pocket, defined by amino acid residues within 6Å of the co-crystalized ligand in PDB ID 7KAC, served as input for SE3-BiLingoMol. The model generated 1.2 million molecular candidates with property distribution displayed in Figure 3B. The generated molecules were subsequently subjected to a multi-stage virtual screening workflow (Figure 3A, more details provided in Supplementary Materials Part 6). The screening workflow primarily consists of two core components: (1) filtering based on molecular conformational validity, binding affinity and chemical validity, followed by (2) clustering based on molecular structure similarity. Specifically, the screening employs multiple submodules, including approximated binding quality measurement using GlideSP score (min-in-place), evaluation of experimental electron density-based ligand-pocket complementarity using ExptGMS^44^, estimation of the occupancy in a subpocket defined by residues L23, G24, and G25, a conformational validity classifier using torsional energy descriptor^28^, drug-likeness assessment using QED and SAS, and additional 2D metrics such as molecular weight, and number of rotatable bonds. Subsequently, the filtered molecules were processed through a clustering module. First, a similarity matrix was obtained by scoring the maximum common substructure of Bemis-Murcko (BM) scaffolds^45^. Subsequently, a spectral clustering was employed to yield the final top 200 molecules. The top 200 candidates (Supplementary Materials Part 7) were then selected for detailed visual inspection, with a particular focus on their predicted interactions with the L23-G24-G25 region and their scaffold novelty.

**Figure 3.**
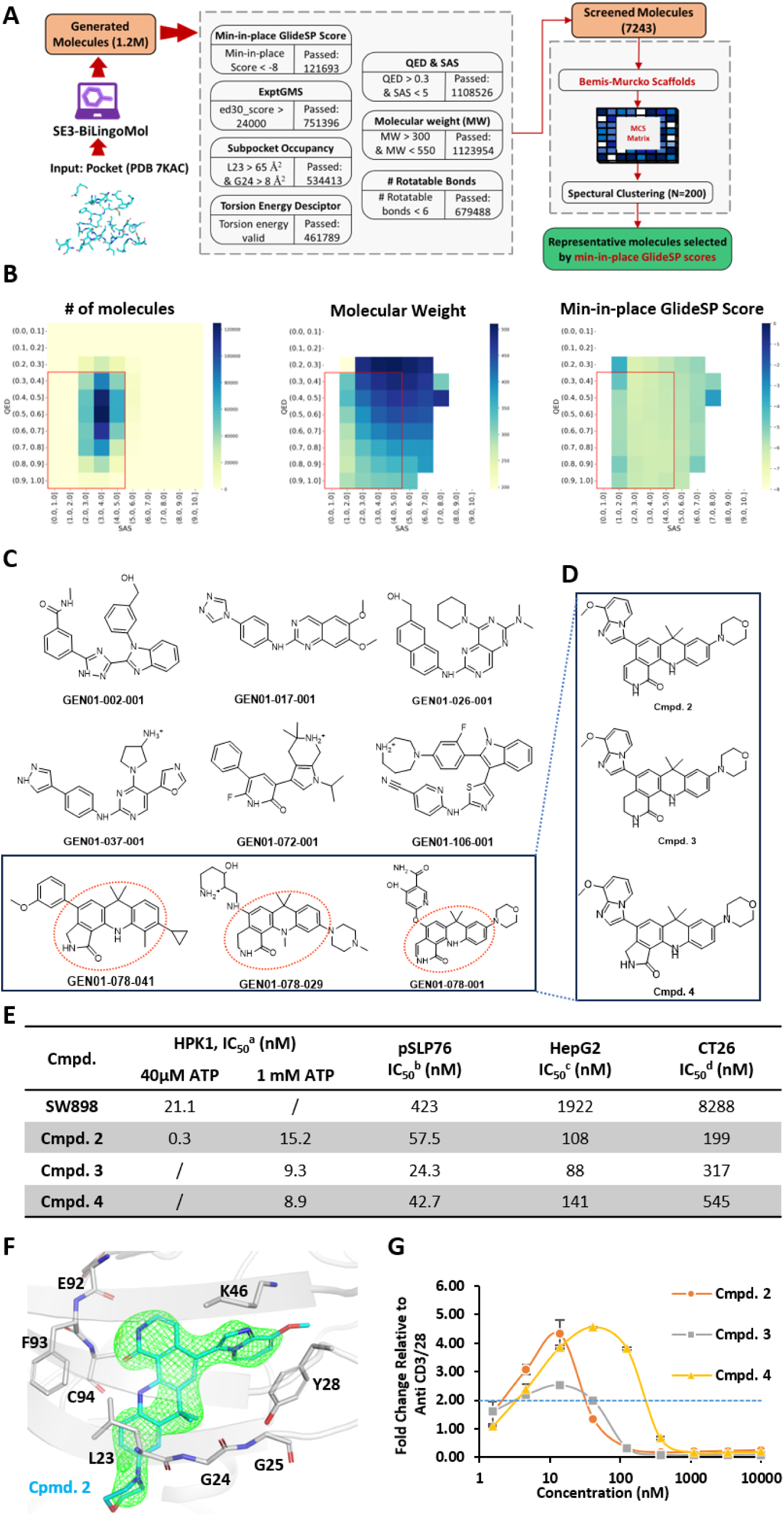
AI-Supported Pocket Structure-based Molecule Design Leading to a Novel Scaffold for HPK1 Hit Inhibitors. (**A**) Schematic diagram illustrating the integrated AI-driven molecular generation and screening pipeline employed for the discovery of novel hit compound scaffolds. In this initial discovery phase, target protein pocket structures served as the input for AI-guided molecule generation. (**B**) Property distribution of the AI-generated molecules. The number of generated molecules, molecular weight (MW), and quality of binding with the pocket (indicated by min-in-place GlideSP scores) are plotted against QED and SAS. The red square indicates drug-like region defined by QED > 0.3 and SAS < 5. (**C**) Representative chemical structures showcasing both classical hinge-binding scaffolds and the novel tetracyclic scaffold series. The tetracyclic cores are explicitly indicated by red dashed circles. (**D**) Structures of compounds synthesized for subsequent biochemical and cellular evaluations. These compounds incorporate the AI-generated tetracyclic cores, while retaining the methoxy imidazopyridine moiety in region A and the morpholine in region C of the reference compound SW898. (**E**) Biochemical and cellular potency of compounds with the tetracyclic cores. ^a^HTRF enzymatic assay. ^b^Jurkat cell assay measuring phosphorylation of SLP76. ^c,d^Characterization of compound toxicity on HepG2 and CT26 cells using the CellTiter-Glo® Luminescent Cell Viability Assay (CTG). (**F**) X-ray crystallographic confirmation of Cmpd. 2 binding within the HPK1 active site (3.2 Å resolution). The F_obs_−F_calc_ omit map, contoured at 3.0 σ (green mesh), confirms the presence of Cmpd. 2 within the HPK1 binding pocket. To generate this unbiased omit map, Cpmd. 2 was excluded from the model by setting its occupancy to zero, followed by several cycles of structural refinement against X-ray diffraction data. (**G**) Bell shaped IL-2 response curves in human primary T cells for Cmpd. 2, 3, and 4. Each data point represents the mean ± SD derived from three independent experimental replicates.

In the initial screening of 200 molecules, while numerous classical hinge-binding scaffolds were identified (Figure 3C, Supplementary Materials Part 8 Figure S5a), a novel tetracyclic scaffold series emerged as particularly noteworthy (Figure 3D, Supplementary Materials Part 8 Figure S5b). Comparing with SW898, this tetracyclic structure exhibits structural similarities to the A region of SW898 (Figure 2A). However, from a prospective standpoint, the conception of fusing a ring with adjacent ring systems to form a novel tetracyclic core was neither readily apparent nor intuitively derivable; rather, it represents an innovative strategy which precisely positions a methyl group within the L23 and G24 region. Although the *in silico* generated molecules incorporating this tetracyclic core displayed diverse substituents at the B and C regions (Figure 3C), Cmpd. 2 combining the novel tetracyclic core with the B and C moieties of SW898 was synthesized and tested (Figure 3D). This compound enables a focused evaluation of the tetracyclic core’s sole impact. As shown in Figure 3E, compared with SW898, Cmpd. 2 demonstrated ∼100-fold improvement of enzymatic potency for HPK1. Furthermore, the X-ray crystal structure of Cmpd. 2 bound to HPK1 was determined at 3.2 Å (Figure 3F, Extended Data Table 2). Cmpd. 2 interacted with HPK1 pocket through non-covalent interactions including hydrogen bonds with the main chain of E92 and C94 in hinge region, π stacking with the side chain of Y28, hydrogen bond with the side chain of K46, and the stacking of methyl from tetracyclic with the amide group of G24 (Extended Data Figure 1A-E). In addition, good alignment between Cmpd. 2 crystal structure and its AI generated counterpart was observed (Extended Data Figure 1F), demonstrating the ability of generating accurate 3D poses of our SE3-BiLingoMol.

However, despite the improvements on activity, Cmpd. 2 still exhibited notable cytotoxicity in HepG2 and CT26 cell lines (Figure 3G). Additionally, single oral dosing in mice resulted in dose-limiting adverse effects, including reduced activity and hypothermia. We hypothesized that the excessive aromaticity and planarity of Cmpd. 2, showing polycyclic aromatic hydrocarbons (PAHs) characteristics which are known to induce cytotoxicity, were responsible for the observed toxicity. To mitigate the aromaticity of the compound, we replaced the tetracyclic moiety in Cmpd. 2 with other similar tetracyclic groups identified from our initial generated series (GEN01-78-029 and GEN01-78-041 shown in Figure 3C), leading to the synthesis of compounds Cmpd. 3 and Cmpd. 4 (Figure 3D). Both derivatives maintained potent HPK1 biochemical activity while demonstrating reduced CT26 cytotoxicity (Figure 3E). We further evaluated their ability to activate IL-2 secretion in human primary T cells. As shown in Figure 3G, both compounds exhibited broader dose windows for IL-2 induction compared to Cmpd. 2, with Cmpd. 4 showing a more expansive 2-fold dose window than Cmpd. 3.

#### 1.2.2. Compound Optimization

Building upon the promising profile of Cmpd. 4, further optimization was pursued through a targeted generation of C-region fragments using the SE3-BiLingoMol model with region A and C fixed, aiming to further diminish overall molecular aromaticity (Figure 4A). This computational effort involved the generation of 0.2 million molecules. A subset of the top 200 molecules (Supplementary Materials Part 9) was then chosen based on a refined screening pipeline (Figure 4A), adapted from the preceding generation round. Specific modifications to this pipeline included the usage of experimental electron density from the Cmpd. 2-HPK1 complex for the ExptGMS assessment, and the mandatory fulfillment of key NCIs involving K46 and Y28. Additionally, the similarity matrix calculation for clustering was changed from previous maximum common substructure (MCS) based scoring to Tanimoto similarity of Morgan fingerprints-based scoring. This alteration was made to emphasize the subtle changes in molecules, as large regions of A and C were retained.

**Figure 4.**
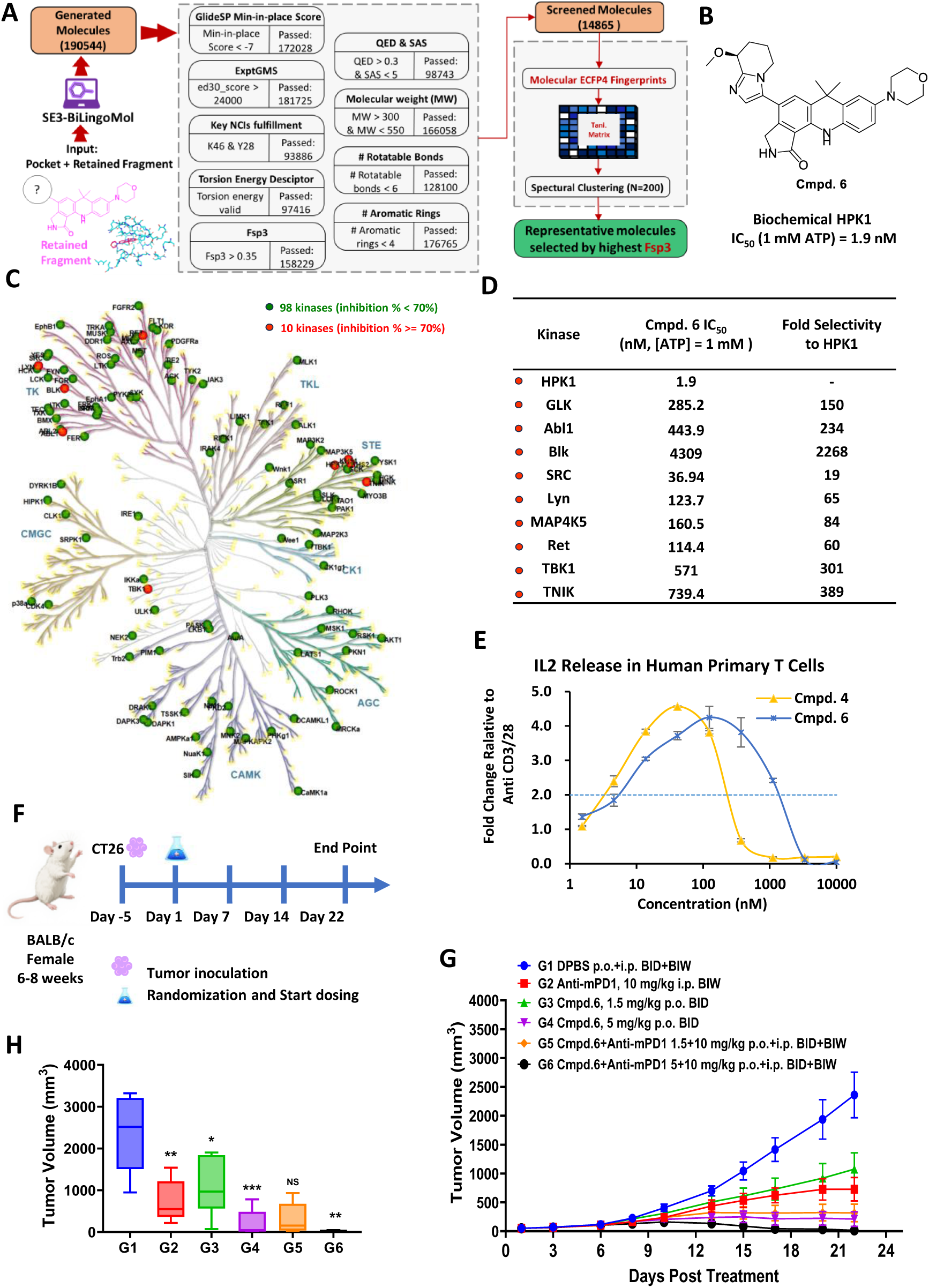
AI-Guided Compound Optimization Leading to an HPK1 Selective Lead Inhibitor with In Vivo Efficacy. (**A**) Schematic diagram of the AI-powered molecule generation and screening pipeline utilized for lead compound optimization. This phase specifically focused on optimizing chemical substituents in region C, while maintaining fixed structures in regions A and B. The target protein pocket structure, complexed with a fragment containing regions A and B derived from Cmpd. 4, served as the input for this round of AI-guided molecule generation. (**B**) Chemical structure of Cmpd. 6, a lead candidate identified through AI-guided optimization. (**C**) Kinome-wide selectivity test for Cmpd. 6. A kinome selectivity analysis was performed involving a total of 108 distinct kinases. Kinase inhibition rates were tested under conditions of 0.03 µM Cmpd. 6 and 40 µM ATP. The 40 µM ATP concentration was chosen, lower than physiological cellular millimolar (mM) levels, to enhance experimental sensitivity and facilitate the detection of potential selectivity liabilities. Kinases exhibiting an inhibition rate of less than 70% are demarcated by green circles, while those demonstrating an inhibition rate equal to or greater than 70% are indicated by red circles. The complete kinome tree, encompassing 518 human protein kinases, was generated using KinMap (http://kinhub.org/kinmap/index.html). (**D**) IC_50_ values and fold selectivity to HPK1 for kinases with ≥70% inhibition. IC_50_ values and the fold selectivity relative to HPK1 are displayed for those kinases exhibiting ≥70% inhibition in the initial kinome screen (as indicated with red circles in Panel C). Fold selectivity was calculated by dividing the IC_50_ of the specific kinase by the IC_50_ of HPK1. For these quantitative IC_50_ determinations, an ATP concentration of 1 mM was used to more accurately assess true selectivity risks, mirroring cellular ATP levels. (**E**) Dose-response for IL-2 Secretion in human primary T cells. Bell-shaped dose-response curve depicting the effect of Cmpd. 6 on IL-2 secretion in human primary T cells. The curve for Cmpd. 4 is included for comparison, illustrating Cmpd. 6’s broad two-fold dose window. Data are presented as mean ±SD from three independent replicates. (**F**) Schematic diagram of the *in vivo* study of Cmpd. 6 in CT26 tumor-bearing mice. (**G**) *In vivo* anti-tumor efficacy in a CT26 mouse model. Tumor volume progression in CT26 tumor-bearing mice treated daily via oral administration with vehicle, Cmpd. 6 (at 1.5 mg/kg and 5 mg/kg), anti-mouse PD-1 (anti-mPD-1) antibody, or their combinations. Values represent mean tumor volume ±SEM (n=6 mice per group). (**H**) Comparison of tumor volume at study endpoint. Tumor volumes on Day 22 of the study were compared. For monotherapy treatment groups (G2, G3, G4), comparisons were made against the vehicle-treated group (G1). For combination treatment groups (G5, G6), comparisons were made against the anti-mPD-1 monotherapy group (G2). A two-sided T-test was employed for statistical analysis, with significance levels indicated as: *p ≤ 0.05, **p ≤ 0.01 (n=6 mice per group).

From this curated set of 200 molecules, by using a molecule dynamics interaction fingerprints (MD-IFPs) analysis^46^ and molecular mechanics-generalized Born surface area (MM/GBSA) calculations^47^ (details provided in Supplementary Materials Part 10), five compounds were selected for synthesis and subsequent comprehensive biochemical and cellular evaluation. As detailed in Extended Data Table 3 and Figure 4B, Cmpd. 6 distinguished itself with optimal biochemical activity, a favorable EC_50_, and the highest maximal fold change in the IL-12 secretion assay in human primary T cells.

Further comprehensive characterization of Cmpd. 6 elucidated its promising pharmacological profile. Cmpd. 6 exhibited good selectivity in a Kinome-wide selectivity test including 108 kinases (Figure 4C), demonstrating notably improved selectivity with >100-fold, >80-fold, >350-fold, and >300-fold against GLK, MAP4K5, TNIK, and TBK1, respectively, which were key off-target kinases problematic for earlier compound SW898 (Figure 4D). It also demonstrated low toxicity in HepG2 and CT26 cell lines (Extended Data Table 4). Importantly, Cmpd. 6 exhibited a broad two-fold dose window for the IL-2 secretion curve in human primary T cells (Figure 4E). Cmpd. 6 also showed good solubility and permeability, and low clearance, which translate to desirable oral bioavailability in pharmacokinetic (PK) studies in both *m*ouse and rat models (Extended Data Table 4). In a mice CT26 tumor model (Figure 4F), once-daily oral administration of Cmpd. 6 showed dose-dependent anti-tumor efficacies (Figure 4G-H). As a monotherapy, Cmpd. 6 achieved tumor growth inhibition (TGI) of 56% and 93% at 1.5 mg/kg and 5 mg/kg, respectively. Combination with a PD-1 antibody further enhanced efficacy, achieving TGIs of 89% and 102%. Safety assessments (Extended Data Table 4) showed that Cmpd. 6 had low risk for CYP inhibition/induction and was Ames test negative. In addition, despite its single-digit nanomolar biochemical and cellular potency, Cmpd. 6’s hERG inhibition occurred only at micromolar concentrations, posing limited safety concern. Collectively, this robust efficacy and safety profile positions Cmpd. 6 as a promising lead compound for HPK1 inhibition in cancer immunotherapy.

## 2. Discussion

Existing molecule generative models, including language models and graph-based models, exhibit substantial limitations in simultaneously addressing 2D drug-likeness, 3D conformational validity, and optimal pocket binding mode. While language models demonstrate proficiency in generating plausible 2D molecular topologies, they inherently lack robust 3D spatial understanding and SE(3)-equivariance. Conversely, graph-based models are adept at capturing 3D atomic relationships and generating valid conformations but often fall short in producing 2D topologies with high drug-likeness. This fundamental dichotomy renders both approaches incomplete for rational drug design. We introduced SE3-BiLingoMol, a novel generative architecture that integrates an E(3)-equivariant Graph Attention Transformer (GATr) with a bidirectional attention mechanism. This design directly overcomes two critical limitations of language models in 3D molecular generation: the inability to handle continuous atomic spatial coordinates with SE(3)-equivariance and the accumulation of conformational errors during autoregressive sampling. Our work significantly advances language models by strengthening their 3D spatial awareness while preserving their established strength in generating chemically plausible 2D structures, thereby transforming them into a comprehensive AI solution for the complex demands of rational drug design. Applied to HPK1 inhibitor design, our model demonstrates a robust AI solution for structure-based drug design (SBDD).

In this discussion, we first elaborate on the significance of our advancements in handling continues geometric properties with SE(3)-equivariance and mitigating conformational errors accumulated during autoregressive generation. Subsequently, we highlight the fundamental differences in 2D chemical rationality between language models and graph-based models, thereby emphasizing how our approach establishes language models as a comprehensive AI solution capable of fully satisfying the multifaceted requirements of SBDD. Building upon this, we discuss the implications of applying SE3-BiLingoMol to support HPK1 inhibitor design, particularly in establishing a paradigm for human-AI interaction. Finally, we discuss the current limitations of our model and directions for future research.

A significant limitation of prior language models for 3D molecule design has been their difficulty in accurately representing the continuous nature of atomic coordinates within 3D protein-ligand complexes. Previous approaches commonly discretized Cartesian coordinates into discrete tokens, treating their prediction as a multi-class classification problem. This discretization often resulted in an inadequate capture of critical binding geometries. In contrast, our model treats atomic coordinates as continuous values and employs a regression-based approach for their prediction. As illustrated in Extended Data Figure 2, the distribution of various bond lengths predicted by models relying on discretized tokens and multi-class classification exhibited a jagged, discontinuous pattern. Conversely, both experimentally determined structures and the predictions from our model yielded smooth, continuous distributions of bond lengths. This observed congruence demonstrates that our method more accurately models the continuous space of molecular geometries. Furthermore, we achieved SE(3)-equivariance in handling these continuous coordinates, which significantly enhances the robustness of inference results. To illustrate this, we applied random rotations and translations on a protein pocket structure six times and subsequently generated molecules using our SE(3)-equivariant model (i.e., SE3-BiLingoMol). These results were then compared with those generated by Lingo3DMol, a language model lacking SE(3)-equivariance. As shown in Extended Data Figure 3, molecules generated by our SE3-BiLingoMol exhibited stable performance across metrics related to their 3D conformational fit within the binding pocket (quantified by pocket occupancy and min-in-place GlideSP score). In contrast, Lingo3DMol’s metrics demonstrated considerable instability, with its performance fluctuating widely for all six randomly rotated inputs.

Beyond addressing 3D special awareness for language models, another critical limitation in these models is the inherent propensity for error accumulation rooted from autoregressive sampling. In such sequential generation, an early error in atom placement or bond formation can propagate and amplify, leading to a cascade of inaccuracies that result in strained bonds, steric clashes, and energetically unfavorable or chemically implausible conformations. SE3-BiLingoMol directly mitigates this pervasive issue through its self-refinement module (Figure 1C). Instead of solely relying on rule-based constraints or post-generation adjustments, this module incorporates a bidirectional attention mechanism that facilitates self-refinement of ligand conformations during the generation process. By enabling global perception of the entire molecular structure in its binding pocket, the module allows for a comprehensive assessment beyond the limited context of preceding tokens. This holistic view facilitates a crucial post-autoregressive local geometry optimization, effectively serving as an intrinsic error-correction layer that significantly alleviates the accumulated errors. Detailed implementation of this module is introduced in Methods Sec, 3.3. We demonstrated the effectiveness of the self-refinement module through an ablation test on this module, keeping all other architectural components consistent. As quantitatively demonstrated in Supplementary Materials Part 11, this mechanism significantly enhances conformation validity (e.g., from 66.6% to 85.2%) and substantially reduces molecular strain energy. For further illustration, we selected PDB 6SCM as a case study (Extended Data Figure 4A). Our SE3-BiLingoMol without self-refinement module was used to generate 9,000 molecules for this target. Across these molecules, we tracked the sampling order of each atom during the autoregressive process (Extended Data Figure 4b), calculating at each corresponding position the probability of having conformational errors (i.e., high-energy atoms as determined by HEAD). This allowed us to visualize the accumulation of 3D coordinate errors during autoregressive sampling. As shown in Extended Data Figure 4C, the probability of having conformational errors exhibited a positive correlation with the atom’s sampling order, a trend that persisted regardless of where the sampling terminated (Extended Data Figure 4D-F). Crucially, after applying the bidirectional attention-driven self-refinement module to these molecules, conformational errors at all positions were notably ameliorated (Extended Data Figure 4C-F). These observations collectively show that our bidirectional attention supported self-refinement model is able to improve the conformation generated in autoregressive sampling process.

As a language model-based architecture, SE3-BiLingoMol circumvents the suboptimal topological drug-likeness issues usually observed in graph-based models. Specifically, we observed a notably high percentage of non-aromatic rings in molecules generated by leading GNN-based models, which is also reported in previous studies^48^. To quantitatively illustrate this prevalent issue, we systematically calculated the percentage of non-aromatic rings across molecules generated by various AI models. As demonstrated in Extended Data Figure 5A, all tested graph-based models—including Pocket2Mol, TargetDiff, PMDM, and MolCraft—consistently generated molecules with a notably higher non-aromatic ring composition compared to their language-model-based counterparts and reference drug molecules from DrugBank^49^. Furthermore, Extended Data Figures 5B and 5C reveal that this undesirable bias is not target-specific but persists across diverse binding targets, indicating a systemic limitation inherent to the model architecture. Extended Data Figures 5D-G showcase examples of molecules generated by these models, frequently exhibiting excessive and often unrealistic fused ring structures. In contrast, SE3-BiLingoMol, along with other language models, produced molecules with a more balanced proportion of aromatic and non-aromatic rings, resulting in fewer topologically flawed or overly complex fused ring structures. This superior topological validity and enhanced drug-likeness in SE3-BiLingoMol stems from its Transformer backbone, which enable a more accurate learning of chemical “grammar” and precise spatial relationships, thus overcoming the intrinsic difficulties GNNs often face in ensuring chemically plausible and drug-like ring systems.

In addition to architectural advancements, our work provides a refined understanding of AI’s role in molecular design by depicting a human-AI interactive workflow in an evidence-based context. Specifically, drug designers tend to seek novel ideas from AI within an evidence-based context, rather than expecting entirely new molecules in a single step without prior knowledge endorsement. The “context” refers to the AI-generated molecule’s scaffold geometry and occupancy within the binding pocket, and it becomes “evidence-based context” when it resonates with validated experimental data. The inclusion of this “evidence-based context” is amenable to Structure-Activity Relationship (SAR) analysis and thereby aligns with drug designers’ established modes of thinking, which improves the human-AI bond. For instance, our SE3-BiLingoMol generated the GEN01-078 quadracyclic series compounds (Figure 3C), which, by structural analogy, can be segmented into three regions (A, B, C) corresponding to the patented molecule SW898 (Figure 2A). This allows for combining a novel ‘A’ region from GEN01-078 with the validated ‘B’ and ‘C’ regions of SW898, thereby facilitating the validation of this new design within a context where the ‘B’ and ‘C’ regions are known to effectively engage their sub-pockets. Extending beyond this series, our model also generated additional compounds incorporating fragments similar to or possessing scaffolds analogous to previously reported active molecules. We synthesized and biochemically evaluated eight of these molecules (Supplementary Materials Part 12), observing that five of them exhibited biochemical inhibitory activity against HPK1, with two achieving sub-50 nM IC_50_ potency. However, their suboptimal engagement with the HPK1 L23-G24-G25 region led to them failing our established filters (Figure 3A) and consequently not entering the top 200 list, while they indeed exhibited lower activity compared to Cmpd. 2.

Next, we will discuss the limitations of SE3-BiLingoMol and potential future directions. While our SE3-BiLingoMol architecture advances the handling of 3D conformations, the generation of energetically favorable conformations at a quantum mechanical (QM) level accuracy remains a frontier. Existing purely data-driven approaches often struggle to capture the intricate details of electron distributions and the delicate coupling between atomic orbitals, where even minute deviations in bond lengths can lead to substantial shifts in the energy landscape. This suggests a crucial future direction: the explicit integration of fundamental physical laws, such as machine learning force fields with QM-level accuracy^50,51^. Additionally, directly leveraging experimental electron density data^44,52,53^ for training represents another powerful avenue to achieve this quantum mechanical precision. These principled approaches are expected to further minimize the strain energy of generated molecules as well as improve their binding mode within the pocket. Beyond the QM-level accuracy of conformation and binding mode in 3D space, the effective exploration of chemical space to generate both novel and synthesizable compounds is also crucial. While SE3-BiLingoMol demonstrates proficiency in generating chemically plausible and drug-like structures, it does not explicitly employ novelty or synthetic accessibility metrics to guide the generation process. Incorporating such metrics could manifest through reinforcement learning^54^ or a conditional generation framework^55^.

In summary, we presented SE3-BiLingoMol as a robust AI solution for SBDD, successfully integrating 2D drug-likeness, 3D conformational validity, and decent pocket binding through its unique GATr architecture. We hope this work can accelerate the discovery of novel therapeutic compounds and inspire future innovations in rational drug design.

## 3. Methods

In this section, we elaborate the detailed methods of SE3-BiLingoMol. We begin with a brief introduction of geometric algebra, followed by an overview of the E(3)-equivariant GATr framework. Because chirality plays an important role in biochemistry system, we showed SE3-BiLingoMol explicitly breaks mirror symmetry to distinguish chiral molecules while preserving SE(3)-equivariant. We then present the key components of our model including the pocket encoder, ligand decoder, self-refinement module and training loss. Next, we illustrate the molecular sampling strategy used during inference.

### 3.1. Geometric Algebra

We will first introduce the concept of geometric algebra (GA) especially for the representation in 3D space, followed by the projective GA.

GA, also known as Clifford algebra, denoted as *g*(V) for 3D space, i.e., V = ℝ^3^, is constructed from an orthogonal basis *e*_1_, *e*_2_, *e*_3_ of the original vector space ℝ^3^, and form a new basis of 2^3^dimensional, i.e., *1,e_1_,e_2_,e_3_,e_12_,e_13_,e_23_,e_123_.*An element spanned by these basis elements is called a multivector. This GA is also equipped with a geometric product, which is a bilinear associative operation, denoted as *xy*, where *x* and *y* are two arbitrary multivectors. However, a representation of this algebra is translation invariant. Thus, instead of using *g*(ℝ^3^), the projective GA, *g*(ℝ^3,0,1^) is constructed to make a representation translation-variant by introducing an additional homogeneous coordinate *e_0_* to the original basis *e_1_,e_2_,e_3_* of Euclidean space, yielding 2^4^-dimensional multivectors *x* ∈ *g*(ℝ^3,0,1^),

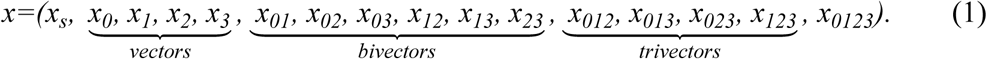

By considering different grades, a multivector can be decomposed into scalars (grade 0), vectors *e*_*i*_ (grade 1), bivectors *e*_*ij*_ (grade 2), trivectors *e*_*ijk*_ (grade 3) and pseudoscalars *e_ijkl_* (grade 4), where*0 ≤ i, j, k, l ≤ 3*. And in such settings, the geometric product for the projective GA satisfies *e*_0_*e*_0_ = 0, *e*_*i*_*e*_*i*_ = 1 for *i = 1, 2, 3*, and *e_i_ e_j_ = –e_j_ e_i_* for *i* ≠ *j*. Notably, a multivector of *g*(ℝ^3,0,1^) is able to represent all the generic objects in 3D space, such as points, lines, planes and symmetry operations, including arbitrary rotations, translations and reflections. For example, representations of scalar features, such as atom types, bond types, and atomic positions are listed in Extended Data Table 5.

### 3.2. Geometric Algebra Transformer

The GATr^32^ is a deep learning model originated from vanilla Transformer^29^, and it was design for handling geometric problems by using a unified architecture. GATr incorporates two key components: (1) the representation of geometric data using projective GA G*_3_*_,*0*,*1*_ and (2) the encoding of E(3) symmetries through equivariant deep learning. To achieve E(3) equivariance, GATr employs several E(3)-equivariant primitives, involving constraining the linear layers of a vanilla Transformer to be equivariant, replacing normalization layers and nonlinearities with equivariant counterparts, enabling the inputs of equivariant multilayer perceptron (MLP) to interact via the geometric product and computing an invariant attention weight between query *q* and key *k* via its inner product. The E(3)-equivariance of GATr’s primitives and the overall architecture is rigorously proven in the original paper^32^.

Like Transformer, GATr utilizes a dot-product attention mechanism, but it operates over multivectors of query, key and value tensors. Because many geometric problems contain non-geometric information, GATr provides auxiliary scalar representations to model these features. These representations mix with scalar components of the multivectors in linear layers and also contribute to attention weight in attention mechanism. In SE3-BiLingoMol, non-geometric features, such as atom types and bond types, are encoded as FSMILES token embeddings and processed as auxiliary scalar representations. The dot-product attention in GATr is formalized as,

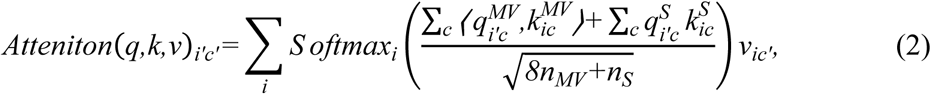

where *q*, *k*, *v* are either multivector-valued (superscript *MV*) or scalar-valued (superscript *S*) query, key and values. The indices *i*, *i′*, *c* and *c′* indicate items and channels of inputs, and *n*_*MV*_ and *n*_*S*_ are the number of multivector and scalar channels, respectively.

### 3.3. SE3-BiLingoMol model architecture

The Figure 1A illustrates the overall architecture of SE3-BiLingoMol. Based on GATr equivariant layers, we define the detailed model components, including the pocket encoder, ligand decoder, ligand head, the self-refinement module and the training loss as follows. The hyperparameters used for SE3-BiLingoMol were listed in Supplementary Materials Part 13.

**Pocket encoder.** The inputs of the encoder involves scalar-valued pocket embeddings 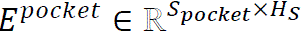 and multivector-valued residue atom positions 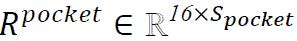, where *S*_*pocket*_, *H*_S_ are the dimensions of the maximum pocket token sequence and hidden size, respectively. The pocket embeddings consist of four parts, namely, token embeddings of residue atoms 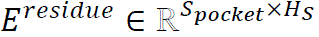, pocket feature embeddings 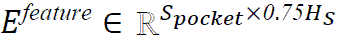, binding interaction embeddings 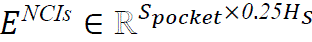 and contact embeddings 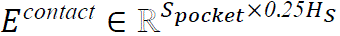. Specifically, binding interaction embeddings *E^NCIs^* and contact embeddings *E^contact^*are firstly summed along its hidden dimension, then followed by a concatenation with pocket feature embeddings *E^feature^*. The pocket embeddings *E*^*pocket*^is then obtained by summing over token embeddings *E*^*residue*^ and the concatenated embeddings along its sequence dimension,

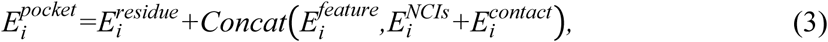

where *i* labels *i*th item of token sequence, and the concatenation of 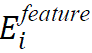 and 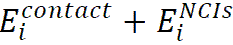 is operated along hidden dimension.

Multivector-valued residue atom positions *R*^*pocket*^is obtained by representing the 3D atomic points as 16-dimensional multivectors as shown in Extended Data Table 5. The pocket encoder, implemented as a GATr module, processes *E*^*pocket*^through the auxiliary scalar channel and *R*^*pocket*^through the multivector channel.

**Ligand decoder.** Similar to the pocket encoder, the ligand encoder is also a GATr module that processes ligand token embeddings 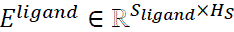 and the corresponding multivector representation of ligand token coordinates 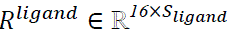. where *S*_*ligand*_is the maximum length of ligand token sequence. Although the decoder maintains E(3) equivariance, we break the E(3) equivariance down to SE(3) equivariance by explicitly distinguishing chiral molecules using ‘@’ and ‘@@’ tokens, similar in the typical SMILES representation.

**Ligand head.** SE3-BiLingoMol incorporates two ligand heads, both implemented as equivariant MLP of GATr primitives. As shown in Figure 1A, the first head is a ligand token head *Head*_*token*_, which predicts the categorical probability distributions of ligand tokens based on the hidden states from the decoder, and the predicted *i*th token is the token with maximum probability,

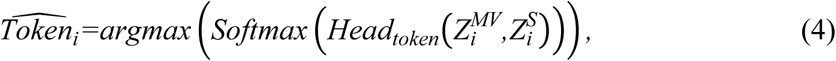

where 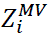 and 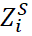 are the *i*th multivector and scalar hiddens states from the decoder. Followed by the second head *Head*_*coords*_, it predicts the multivector representation of the *i*th token 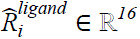 using the *i*th hidden states and the embeddings of the predicted token 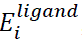,

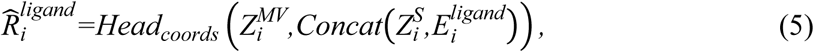

where the embeddings of *i*th token is appended to *i*th scalar embeddings along channel dimension. Finally, the 3D coordinates of *i*th token, *x*_*i*_, *y*_*i*_, *Z*_*i*_, are extracted directly from the predicted multivector representation 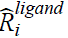,

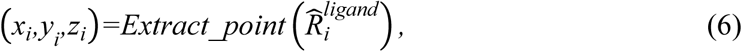

The extraction operator is simply an inverse transformation from 16-dimension space to 3D space as listed in Extended Data Table 5.

**Self-refinement module.** This module adopts the same architecture as the ligand decoder but employs a bidirectional attention mechanism, and it also comprises an additional ligand coordinate head to generate the refined molecular conformation. The bidirectional mechanism allows each ligand token to interact with all other ligand tokens and pocket tokens, so that the molecule structure can be relaxed after the perception of the global context, as shown in Figure 1C. The outputs of this decoder is further used by the ligand coordinate head to predict the relaxed molecule positions.

**Training loss.** During training, the total loss is a weighted sum of ligand token loss *L*_*tokens*_, ligand coordinate loss *L*_*coords*_, and three auxiliary loss functions including ligand bond length loss *L*_*bonds*_, bond angle loss *L*_*angles*_ and dihedral angle loss *L*_*diℎedrals*_,

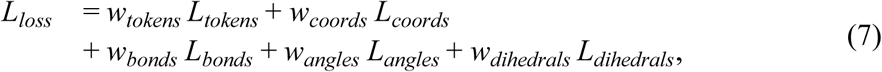

where *w*_*tokens*_, *w*_*coords*_, *w*_*bonds*_, *w*_*angles*_ and *w*_*diℎedrals*_ are the weights of the corresponding loss terms. Specifically, during training, the molecular topology of ground-truth molecules were recorded, and thus, these auxiliary loss terms can be computed based on the topology. Then, the cross entropy loss is used for each *L*_*tokens*_ and SmoothL1Loss is used for others. For the auxiliary losses, these that can form a bond or define a dihedral were only considered, and with the periodic boundary condition considered for *L*_*diℎedrals*_.

### 3.4. Sampling strategy

As illustrated in Figure 1B. at the beginning (T=0), the input binding pocket is prepared by selecting residues within 6Å of a reference ligand (Step 1). Simultaneously, a residue atom position near the pocket center is randomly initialized as the coordinate of the ‘[START]’ token (Step 2). Optionally, a molecular substructure can be provided as an initial seed fragment, whose FSMILES tokens and spatial coordinates will be appended to “[START]” token (Step 3, optional). Then, the pocket and ligand embeddings are processed by the encoder and decoder (Step 4 and 5). Next, the ligand head autoregressively predicts the next token and its position based on previous token and the pocket contextual hidden states (Step 6). Sampling terminates when an “END” token is predicted or the maximum ligand sequence length, *S*_*max*_, is reached.

### 3.5. Protein-ligand complex dataset preparation

Invalid ligands and common buffer molecules were removed. To ensure each entry represented a unique and independent geometric unit, only the asymmetric unit of the crystal structure was retained for each selected complex, rather than its complete biological assembly. Before further analysis, each protein-ligand complex underwent rigorous pre-processing using Schrödinger’s PrepWizard to correct bond orders, assign proper protonation states, and add missing hydrogen atoms, thereby ensuring the structures were suitable for downstream computational workflows. Subsequently, for each prepared complex, the Open Drug Discovery Toolkit^56^ was utilized to identify non-covalent interactions (NCIs) between the ligand and its binding protein. Finally, instead of using entire protein structures, the binding pockets were extracted, defined as all residues within a 6Å distance of the ligand molecule.

### 3.6. Expression and purification of human HPK1 kinase domain

The gene encoding human HPK1 kinase domain (Asp2-Pro298) was inserted into a pFastBac vector fused with an 8xHis tag followed by an HRV 3C protease site at the N-terminus. The inactive mutation S171A was generated by standard site-directed mutagenesis. The plasmids were transformed into DH10bac cells and recombinant bacmids were produced, which were then transfected into Spodoptera frugiperda Sf9 cells using Cellfectin-II (Invitrogen). The baculoviruses were amplified three times in Sf9 cells. P3 viruses were used to transfect Hi5 insect cells at a ratio of 1:50 (v/v) when cells grow to 2.5 x106 cells/ml. After culturing in ESF921 medium for 3 days, the cells were collected by centrifugation and were stored at –80°C.

The cell pellets were resuspended in buffer A (50 mM HEPES, 300 mM NaCl, 0.5 mM TCEP, and 10% glycerol) supplemented with protease inhibitors. After cell breaking by ultrasonication, the cell lysate was centrifugated for 30 min at 45000 rpm and supernatant was incubated with Ni-NTA resins. The resins were washed by 50 mL buffer A supplemented with 20 mM imidazole, followed a second washing with 35 mM imidazole. The HPK1 protein samples were eluted by 20 mL elution buffer (20 mM HEPES, 300 mM NaCl, 0.5 mM TCEP, and 300 mM imidazole). The His tag was cleaved by PreScission protease (1:50 w/w ratio) digestion for 2 h on ice. Subsequently, the samples were concentrated and loaded onto a Superdex 200 Increase10/300 GL column (Cytiva) equilibrated in buffer containing 20 mM HEPES (pH 7.5), 300 mM NaCl and 0.5 mM TCEP. The peak fractions were collected, concentrated to 5 mg/mL and stored at –80°C. Peak fractions were pooled and concentrated to 20 mg/mL for crystallization. Aliquots of the concentrated HPK1 kinase samples were stored at –80°C.

### 3.7. Crystallization, Data Collection and Structure Determination

Prior to crystal screening, the HPK1 kinase samples were incubated with 1 mM Cmpd. 2 for 1 h on ice. Initial needle-shaped crystals were observed in Crystal Screen Kit G6 (0.1 M HEPES pH 7.5, 10% w/v Polyethylene glycol 6,000, 5% v/v (+/-)-2-Methyl-2,4-pentanediol). After condition optimization, 1μL protein solution was mixed with 1μL reservoir solution (0.1 M HEPES pH 6.9, 8% w/v Polyethylene glycol 6,000, 5% v/v (+/-)-2-Methyl-2,4-pentanediol, 10 mM GSH/GSSG) for each drop, and large rod-shaped crystals were obtained after 3-5 days at 18°C. Crystals were transferred into cryoprotectant containing mother reservoir solution supplemented with 25% ethylene glycol, then flash frozen in liquid nitrogen.

X-ray diffraction data of HPK1-Cmpd. 2 crystals were collected at beamline BL17U1 of the Shanghai Synchrotron Radiation Facility (SSRF). X-ray diffraction data were processed using the program HKL2000. The initial phase was solved by molecular replacement (MR) with the HPK1 kinase structure (PDB code: 6CQE) as a reference model using the program Phaser in PHENIX. Several rounds of manual model building and refinements were performed iteratively using COOT and PHENIX refinement. Statistics for data collection and processing are shown in Extended Data Table 1.

### 3.8. Compound Synthesis

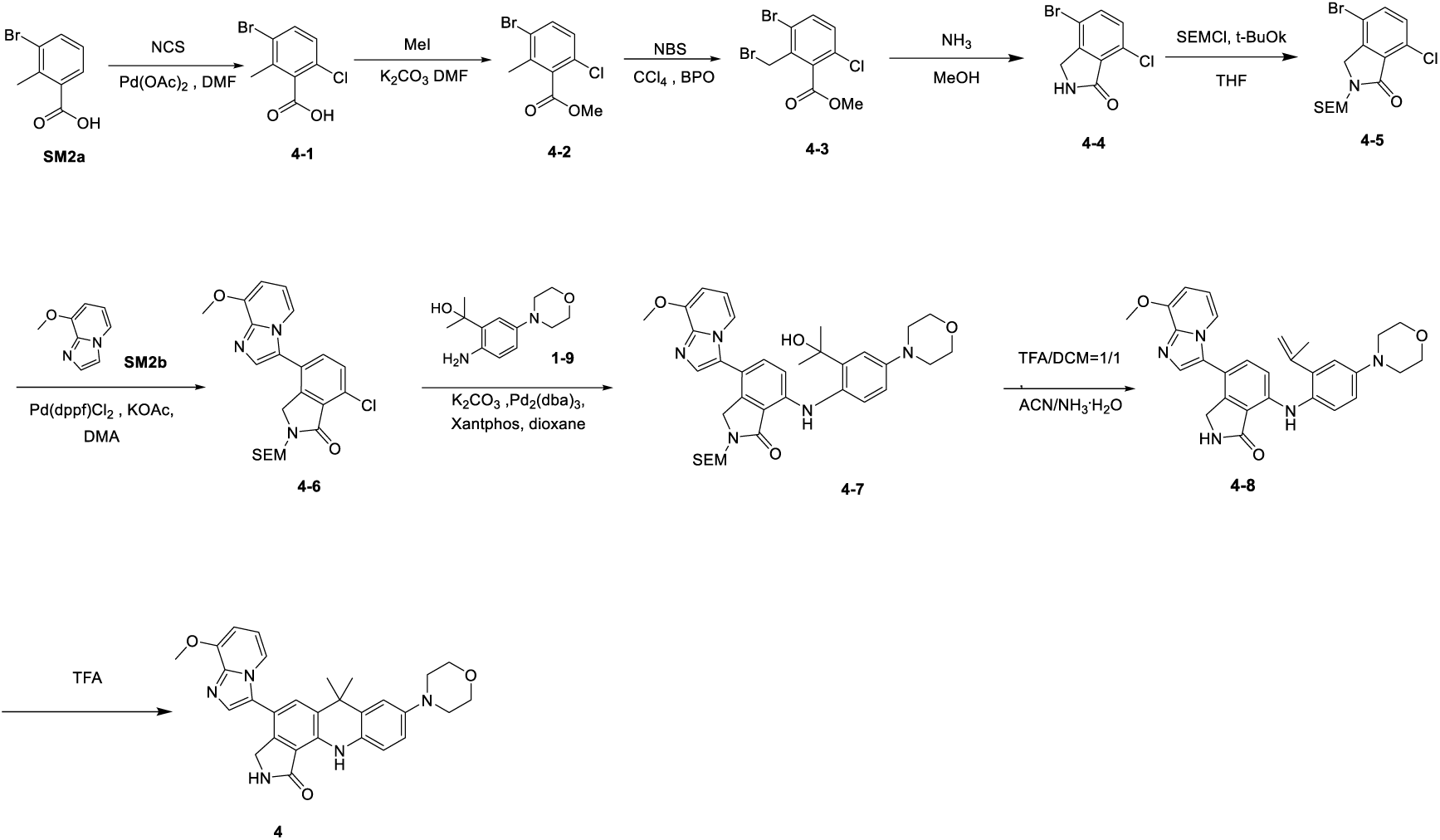

#### Synthesis of compound 4

#### Synthesis of intermediate 4-1

Compound SM2a (200 g) was dissolved in N, N-dimethylformamide (1 L), and NCS (150 g) and palladium (II) acetate (20.9 g) were added. The above mixture was reacted at 110°C for 16 hours. The reaction solution was poured into water (2 L), and the mixture was extracted with ethyl acetate (2 L × 3). The organic phases were combined, dried over anhydrous sodium sulfate, filtered, and rotary evaporated to dryness. The resulting crude product was purified by column chromatography (silica, petroleum ether/ethyl acetate = 1/0 to 0/1) to give yellow solid compound 4-1 (118 g, yield: 50.9%). LCMS (ESI) m/z: 249/251.

#### Synthesis of intermediate 4-2

Compound 4-1 (88 g) was dissolved in N, N-dimethylformamide (620 mL), and iodomethane (55 g) was added. The above solution was reacted at 110℃ for 16 hours. The reaction solution was cooled to room temperature and poured into water (1.8 L). The mixture was extracted with ethyl acetate (1.0 L × 3). The combined organic phase was dried over anhydrous sodium sulfate, filtered, and rotary evaporated to dryness to give yellow oily compound 4-2 (86 g) as a crude product, which was directly used in the next reaction. LCMS (ESI) m/z: 263/265.

#### Synthesis of intermediate 4-3

Compound **4-2** (86 g) was dissolved in carbon tetrachloride (860 mL), and N-Bromosuccinimide (63.8 g) and benzoyl peroxide (3.95 g) were added. The above mixture was reacted at 80°C for 16 hours. The reaction solution was cooled to room temperature and filtered. The filtrate was rotary evaporated to dryness to give yellow oily compound **4-3** (106 g) as a crude product, which was directly used in the next reaction. LCMS (ESI) m/z: 341/343.

#### Synthesis of intermediate 4-4

Compound **4-3** (106 g) was dissolved in methanol (530 mL), and a solution of ammonia in methanol (530 mL) was added under an ice-water bath. The above reaction solution was reacted at room temperature for 4 hours. The reaction solution was filtered, and the filter cake was dried to give off-white solid compound **4-4** (62 g) as a crude product, which was directly used in the next reaction. LCMS (ESI) m/z: 246/248.

#### Synthesis of intermediate 4-5

Compound **4-4** (62 g) was dissolved in anhydrous tetrahydrofuran (1 L), and potassium tert-butoxide (56.5 g) and 2-(trimethylsilyl)ethoxy methyl chloride (125.8 g) were added under an ice-water bath. The above mixture was reacted at room temperature for 16 hours. The reaction solution was added to water (1 L), and the mixture was extracted with ethyl acetate (1 L x 3). The combined organic phase was dried over anhydrous sodium sulfate, filtered, and rotary evaporated to dryness. The resulting crude product was purified by column chromatography (silica, petroleum ether/ethyl acetate = 1/5 to 1/1) to give yellow oily compound **4-5** (43 g). LCMS (ESI) m/z: 376/378. ^1^H NMR (400 MHz, CDCl_3_) δ ppm: 7.61 (d, *J* = 8.4 Hz, 1H), 7.33 (d, *J* = 8.4 Hz, 1H), 5.06 (s, 2H), 4.38 (s, 2H), 3.63 – 3.56 (m, 2H), 1.01 – 0.91 (m, 2H), 0.01 (s, 9H).

#### Synthesis of intermediate 4-6

Compound **4-5** (380 mg) and compound **SM2b** (179 mg) were dissolved in N, N-dimethylacetamide (2 mL), and potassium acetate (198 mg) was added. The reaction system was purged with nitrogen three times, and then [1,1’-Bis(diphenylphosphino)ferrocene] palladium (II) dichloride (73.9 mg) was added. The above mixture was stirred at 110°C for 3 hours under nitrogen. The reaction solution was cooled to room temperature and filtered, and the filtrate was rotary evaporated to dryness. The resulting crude product was purified by column chromatography (silica, petroleum ether/ethyl acetate = 1/5 to 0/1) to give brown solid compound **4-6** (260 mg, yield: 58%). LCMS (ESI) m/z: 445.

#### Synthesis of intermediate 4-7

Compound **4-6** (120 mg) and compound **1-9** (63.8 mg) were dissolved in dioxane (5 mL). Potassium carbonate (114 mg) and 4,5-Bis(diphenylphosphino)-9,9-dimethylxanthene (31.2 mg) were added. The reaction system was purged with nitrogen three times, and then Tris(dibenzylideneacetone)dipalladium (0) (24.7 mg) was added. The above mixture was stirred at 100°C for 12 hours under nitrogen. The reaction solution was cooled to room temperature and filtered, and the filtrate was rotary evaporated to dryness. The resulting crude product was purified by column chromatography (silica, petroleum ether/ethyl acetate = 1/3 to 0/1) to give brown solid compound **4-7** (85 mg, yield: 49%). LCMS (ESI) m/z: 644.

#### Synthesis of intermediate 4-8

Compound **10-7** (85 mg) was dissolved in dichloromethane (1 mL), and trifluoroacetic acid (1 mL) was added dropwise. The above solution was stirred at room temperature for 2 hours. The reaction solution was concentrated under reduced pressure to give a brown oily compound. The brown oily compound was dissolved in acetonitrile (1 mL). Ammonia solution (1 mL) was added, and the mixture was stirred at room temperature for another 10 hours. The mixture was rotary evaporated to dryness and purified by high performance liquid chromatography (chromatography column: Waters XBridge BEH C18 100*30 mm*10 um; mobile phase: [water (ammonium bicarbonate) – acetonitrile]; B%: 30% – 60%, 8 min) to give yellow solid compound **4-8** (36 mg, yield: 57%). LCMS (ESI) m/z: 496.

#### Synthesis of compound 4

Compound **4-8** (34 mg) was added to trifluoroacetic acid (3 mL). The above solution was stirred at 80°C for 12 hours. The reaction solution was rotary evaporated to dryness, and purified by high performance liquid chromatography (chromatography column: Phenomenex Luna C18 75*30mm*3um; mobile phase: [water (formic acid) – acetonitrile]; B%: 1% – 40%, 8 min) to give yellow solid compound **4** (4.4 mg, yield: 12.9%). LCMS (ESI) m/z: 496. ^1^H NMR (400 MHz, DMSO-*d_6_*) δ ppm: 8.91 (s, 1H), 8.57 (s, 1H), 7.84 (d, *J* = 6.8 Hz, 1H), 7.68 (d, *J* = 6.4 Hz, 2H), 7.05 (d, *J* = 2.4 Hz, 1H), 6.92 (d, *J* = 8.8 Hz, 1H), 6.85 (t, *J* = 7.2 Hz, 1H), 6.77 (dd, *J* = 2.4, 8.8 Hz, 1H), 6.71 (d, *J* = 7.2 Hz, 1H), 4.28 (s, 2H), 3.96 (s, 3H), 3.76 – 3.73 (m, 4H), 3.05 – 3.02 (m, 4H), 1.61 (s, 6H).

Other compounds, including Cmpd. 2, 3, 5, 6, 7, 8, and 9, were synthesized using methods highly similar to those described above.

## 4. Declarations

### 4.1. Data and code availability

The DUD-E dataset is a publicly available via link https://dude.docking.org/ Our source code and partial training data have been deposited in Ocean Code and Figshare, respectively, for peer review. These resources will be made publicly accessible following the publication of the manuscript.

## Acknowledgements

This work was supported by the Beijing Municipal Science and Technology Commission (grant no. Z241100007724005 received by B.H.). This study was also funded by the National Key R&D Program of China (grant no. 2022YFF1203004 received by B.H.).

## 4.2. Author contributions

B.H. conceived the study. W.Z. provided instructions for artificial intelligence modelling. H.W provided instructions for compound evaluation. B.H. and Z.L. provided instructions on evaluation framework. D.J. provided instructions for structural biology study. B.X. and H.W. build the AI model. G.S synthesize and evaluated the compounds. B.Z prepared the training data. Y. W., R.M, Y.P. and F.Z. supported the model evaluation. Y. G. and Y.W. supported the compound evaluation.

## 4.3. Competing interests

The authors declare no competing interests.

**For 2D Molecular Quality**, assessment is generally performed by evaluating a Quantitative Estimate of Drug-likeness (**QED**)^41^ score and a Synthetic Accessibility Score (**SAS**)^42^. The **% Druglike Molecules** metric represents the percentage of generated molecules exhibiting a **QED** score greater than 0.3 and a **SAS** less than 5. More specific metrics such as **% Aromatic rings** and **% Non-aromatic rings** are also provided when the general metric of **% Druglike Molecules** is not sufficiently distinguishing. The **% Aromatic rings** metric estimates the average ratio of aromatic rings in the generated molecules. Specifically, for each molecule, the atomic masses of all atoms in aromatic rings (determined using RDKit) are summed, and the aromatic ring ratio is defined as the ratio of the atomic masses of aromatic-ring atoms to its total molecular weight. The **% Non-aromatic rings** metric estimates the average ratio of non-aromatic rings in the generated molecules. Specifically, for each molecule, the atomic masses of all atoms in non-aromatic rings (determined using RDKit) are summed, and the non-aromatic ring ratio is defined as the ratio of the atomic masses of atoms of non-aromatic rings to its total molecular weight. For drug like molecules (QED>0.3 & SAS<5), **Chemical Diversity** is also provided. It measured as 1 minus the average Tanimoto similarity between unique pairs of generated molecules (ECFP4 fingerprints).

**For 3D Conformation Quality regarding intrinsic ligand conformations**, we report the **% Conformation Validity**, which is the percentage of molecules that successfully pass local abnormal conformation checks using HEAD^28^. We also present **Strain Energy 25%, 50%, and 75%** to reflect the distribution of molecules’ strain energies (in kcal/mol) calculated using PoseCheck^43^.

**For 3D Conformation Quality regarding Ligand-Pocket Binding Mode**, we report the **Mean Pocket Occupancy,** representing the weighted average occupancy similarity between generated ligands and the co-crystallized reference ligand associated with the PDB ID in DUD-E, evaluated across multiple targets. To compute this score, the solvent-accessible surface area (SASA) difference between the unbound pocket (defined as all residues within 6 Å from its cocrystal ligand) and the ligand-bound complex is calculated for both the co-crystal ligand and each generated ligand. The occupancy of a given ligand is then defined as the ratio of its SASA difference to that of the co-crystal ligand, with values exceeding 1 clipped to 1. The **Mean Min-in-place Score**, quantifying binding quality using the OPLS3e force-field via GlideSP^6^ in “mininplace” mode. The **Mean Redocking Score**, calculated by redocking generated molecules into the pocket via GlideSP in “docking” mode, serves purely as a reference. This is because it involves resampling molecular conformation, not using the generated conformation, thus misaligning with 3D molecule generation objectives. It provides context to understand the **% Min-in-place < Redocking Score**, which indicates the percentage of molecules with a lower (better) min-in-place score than their re-docking score, highlighting the superiority of AI-generated conformations over force-field sampled ones. **% IMP**^57^ is the percentage of generated molecules with a lower min-in-place GlideSP score than that of their active ligand counterparts’ best (lowest) GlideSP docking score on that target. The **PLIF Recovery Rate**^58^ represents the proportion of molecules that replicate the same non-covalent interactions (NCIs) – including hydrogen bond acceptor/donor, halogen bond acceptor/donor, π-π stacking, cation-π, π-cation, anionic, and cationic interactions, as calculated by ProLIF^59^ – as those of the co-crystalized ligands with specific protein residues.

**For Active Reproduction Capability**, the **% ECFP_TS > 0.5** metric^30^ represents the percentage of targets where at least one generated molecule exhibits a Tanimoto similarity (based on ECFP4 fingerprints) greater than 0.5 with a known active compound for that target. To ensure robust selection of active compounds, we docked all active compounds into the pocket indicated by the co-crystallized ligand, and only those with a docking score < –5 were considered. Full details on metric calculation and experimental setup are provided in Supplementary Materials Part 2.

