## Extended Data Figures and Tables for "Pocket-based molecule generation with an SE(3)-equivariant language model leads to a potent and selective HPK1 inhibitor with *in vivo* efficacy"

### Supplementary Information

Bin Xi[1,2†], Han Wang[1,2†], Guanglong Sun[2†], Bowen Zhang[3†], Ruihan Mao[2], Yuyang Ge[2], Yang Wang[2], Jiangtao Zhang[3], Yiting Pan[2], Feng Zhou[2], Yuji Wang[4], Zhenming Liu[1]*, Daohua Jiang[3]*, Huting Wang[2]*, Wenbiao Zhou[2]* and Bo Huang[1,2,4]*

[1]State Key Laboratory of Natural and Biomimetic Drugs, School of Pharmaceutical Sciences, Peking University, Beijing, China.

[2]Beijing StoneWise Technology Co Ltd., Haidian Street #15, Beijing, 100080, China.

[3]Laboratory of Soft Matter Physics, Institute of Physics, Chinese Academy of Sciences, Beijing, China.

[4]Department of Medicinal Chemistry, College of Pharmaceutical Sciences of Capital Medical University, Beijing 100069, P. R. China.

[†]These authors contributed equally to this work.

;


**Extended Data Table 1.** Inference time cost of generating 500 valid molecules on randomly selected five targets for Pocket2Mol, TargetDiff, Lingo3DMol, PocketFlow, PMDM, MolCraft, SE3-BiLingoMol. The experiment was conducted on NVIDIA Tesla V100.

| | **Pocket2Mol** | **TargetDiff** | **Lingo3DMol** | **PocketFlow** | **PMDM** | **MolCraft** | **SE3-BiLingoMol (Ours)** |
|---|---|---|---|---|---|---|---|
| **Time cost (s, ↓)** | **796±366** | **12329±3860** | **2799±809** | **1046±64** | **4390±2228** | **1227±369** | **87±22** |

**Extended Data Table 2. Crystallography Information of HPK1-Cmpd. 2 Complex**

| | HPK1-Cmpd.2 (PDB: 9WD3) |
|---|---|
| Data collection | |
| Space group | P21 |
| Unit Cell | a=116.6 Å, b=99.6 Å, c=117.0 Å<br>α=γ= 90°, β=90.4° |
| Resolution | 3.23 Å (3.40-3.23) |
| Total measured reflections | 94,927 |
| Completeness (%) | 92.0 (99.8) |
| Redundancy | 3.0 (3.2) |
| I/σ | 6.4 (2.6) |
| $R_{merge}$ | 0.117 (0.475) |
| CC(1/2) | 0.984 (0.797) |
| Resolution range | 22.71 - 3.23 Å |
| Refinement | |
| Initial model | PDB code: 6CQE |
| R, $R_{free}$ | 25.0%, 30.2% |
| Model composition | |
| Non-hydrogen atoms | 18014 |
| Protein atoms | 17643 |
| Ligand atoms | 380 |
| Average B factors ($Å^2$) | |
| Protein | 80.9 |
| Ligand | 96.0 |
| R.m.s. deviations | |
| Bond lengths (Å) | 0.013 |
| Bond angles (°) | 1.099 |
| Validation | |
| MolProbity score | 2.30 |
| Poor rotamers (%) | 0.90 |
| Ramachandran plot | |
| Favored (%) | 90.84 |
| Allowed (%) | 7.72 |
| Disallowed (%) | 1.44 |

**Extended Data Table 3. Biochemical and Cellular Evaluation of Compounds Leading to the Selection of Cmpd. 6**

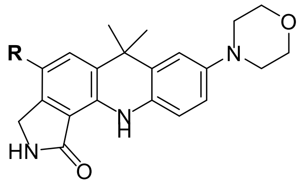

| Cmpd. | R | HPK1, $IC_{50}$ (nM) 1 mM ATP | IL2 PBMC, $EC_{50}$ (nM) | IL2 PBMC, max fold change |
|---|---|---|---|---|
| **5** | 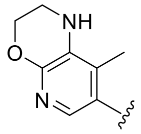 | 3.9 | 6.4 | 2.9 |
| **6** | 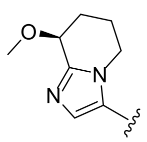 | 1.9 | 9.5 | 4.2 |
| **7** | 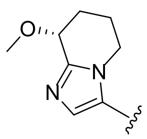 | 4.2 | 5.9 | 2.3 |
| **8** | 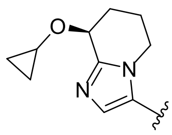 | 5.4 | 15.4 | 3.3 |
| **9** | 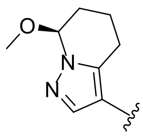 | 11.0 | 24.8 | 3.7 |

**Extended Data Table 4. Cellular Activity, Pharmacokinetic (PK) and Safety Profile of Cmpd. 6.**

| Compound ID | Cmpd. 6 |
|---|---|
| CTG assay, $IC_{50}$ (nM) for HepG2, CT26 | 3286, 7796 |
| IL-2 increase in human primary T cells, $EC_{50}$ (nM), max fold change, 2-fold dose window | 9.5, 4.16, ~380 |
| Solubility (KS, pH=6.5, 7.4) (µM) | 62.6, 56.3 |
| Permeability (Caco-2, A->B; B->A), Mean Papp ($10^{-6}$ cm/s), Efflux Ratio | 5.64, 15.33, 2.72 |
| Hepatocyte Clint (mL/min/kg), $t_{1/2}$ (min) (human, mouse, rat, dog) | 1.56, 2263 (h); 101, 161 (m); 27.5, 236 (r); 46.5, 205 (d) |
| [a]PPB% (human, mouse, rat, dog) | 95.4, 97.7, 96.14, 98.88 |
| CYP1A2, 2C9, 2C19, 2D6, 3A4/5 inhibition, (10 µM) | 9%, 24.5%, 53.6%, 20.7%, 60.5% |
| CYP3A4 [b]TDI, without / with preincubation, $IC_{50}$ (µM) | 5.37 / 5.97 |
| [c]PXR activation as % of [d]positive control (1 uM, 10 uM) | 2%, 2% |
| IV. Mouse PK (1mg/kg): $t_{1/2}$ (hour), Cl (mL/min/kg), $AUC_{0\text{-last}}$ (h*ng/mL), $Vd_{ss}$ (L/kg) | 2.32, 6.72, 2528, 1.03 |
| IV. Rat PK (1mg/kg): $t_{1/2}$ (hour), Cl (mL/min/kg), $AUC_{0\text{-last}}$ (h.ng/mL), $Vd_{ss}$ (L/kg) | 1.49, 14, 1187, 1.63 |
| PO. Mouse PK (3 mg/kg): $t_{1/2}$ (hour), $C_{max}$ (ng/mL), $AUC_{0\text{-last}}$ (h.ng/mL), F% | 2.24, 1982, 7604, 97.7% |
| PO. Rat PK (3 mg/kg): $t_{1/2}$ (hour), $C_{max}$ (ng/mL), $AUC_{0\text{-last}}$ (h.ng/mL), F% | 3.83, 711, 4060, 113% |
| hERG inhibition, (10 µM) | 74.5% (10 µM), $IC_{50}$ = 5.16 µM |

[a]PPB = plasma protein binding (%). [b]TDI = Time-dependent Inhibition. [c]PXR= Pregnane X Receptor. [d]positive control= Rifampicin (10 µM)

**Extended Data Table 5. Representations of scalar features, 3D points and symmetry operations using the projective geometric algebra $G(R^{3,0,1})$.** The non-zero component coefficients of the 16-dimensional multivector x are listed, and the a generic multivector can be expressed as in Eq. 1

| Geometric objects/ symmetry operations | Multivector | mapping |
|---|---|---|
| Scalar $s \in R$ | $x_s$ | $= s$ |
| Point $p \in R^3$ | $(x_{012}, x_{013}, x_{023}, x_{123})$ | $= (p, 1)$ |
| Translation $t \in R^3$ | $(x_s, x_{01}, x_{02}, x_{03})$ | $= (1, t/2)$ |
| Rotation expressed as quaternion $q \in R^4$ | $(x_s, x_{01}, x_{02}, x_{03})$ | $= q$ |
| Point reflection through $p \in R^3$ | $(x_{012}, x_{013}, x_{023}, x_{123})$ | $= (p, 1)$ |
| Reflection through plane w/ normal $n \in R^3$ , origin shift $d \in R$ | $(x_0, x_1, x_2, x_3)$ | $= (d, n)$ |

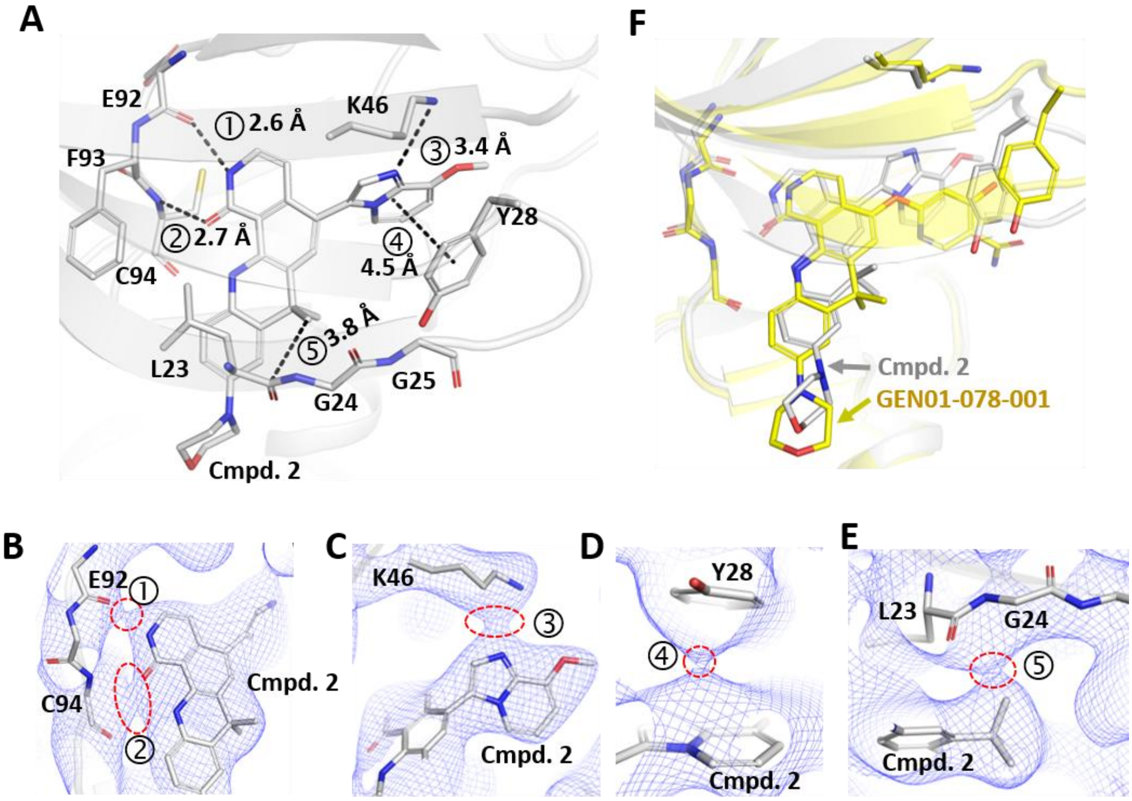

**Extended Data Figure 1. Crystal Structure of Cmpd. 2 in Complex with HPK1 and Comparison with the AI-Generated Counterpart.**

**(A) Binding mode of Cmpd. 2 in HPK1 pocket.** Five non-covalent interactions (NCIs) are numerically labeled, with corresponding interatomic distances indicated.

**(B-E) Bond critical point (BCP) analysis elucidating key non-covalent interactions.** BCPs, indicated by red dashed-line circles in the $2F_{obs}-F_{calc}$ electron density maps, provide atomic-level evidence for NCIs. Contour levels were 0.5 σ for **(B)** and **(C)**, -0.3 σ for **(D)**, and 0.0 σ for **(E)**.

**(F) Superposition of experimentally determined Cmpd. 2 and its AI-generated counterpart.** The experimentally determined HPK1-Cpd. 2 complex (white) is superimposed with the AI-generated molecule (yellow), GEN01-078-001 (Figure 3C), which shares the same tetracyclic core. The AI molecule was generated using the pocket structure of PDB entry 7KAC. The experimental complex's protein backbone was aligned with that of the 7KAC protein.

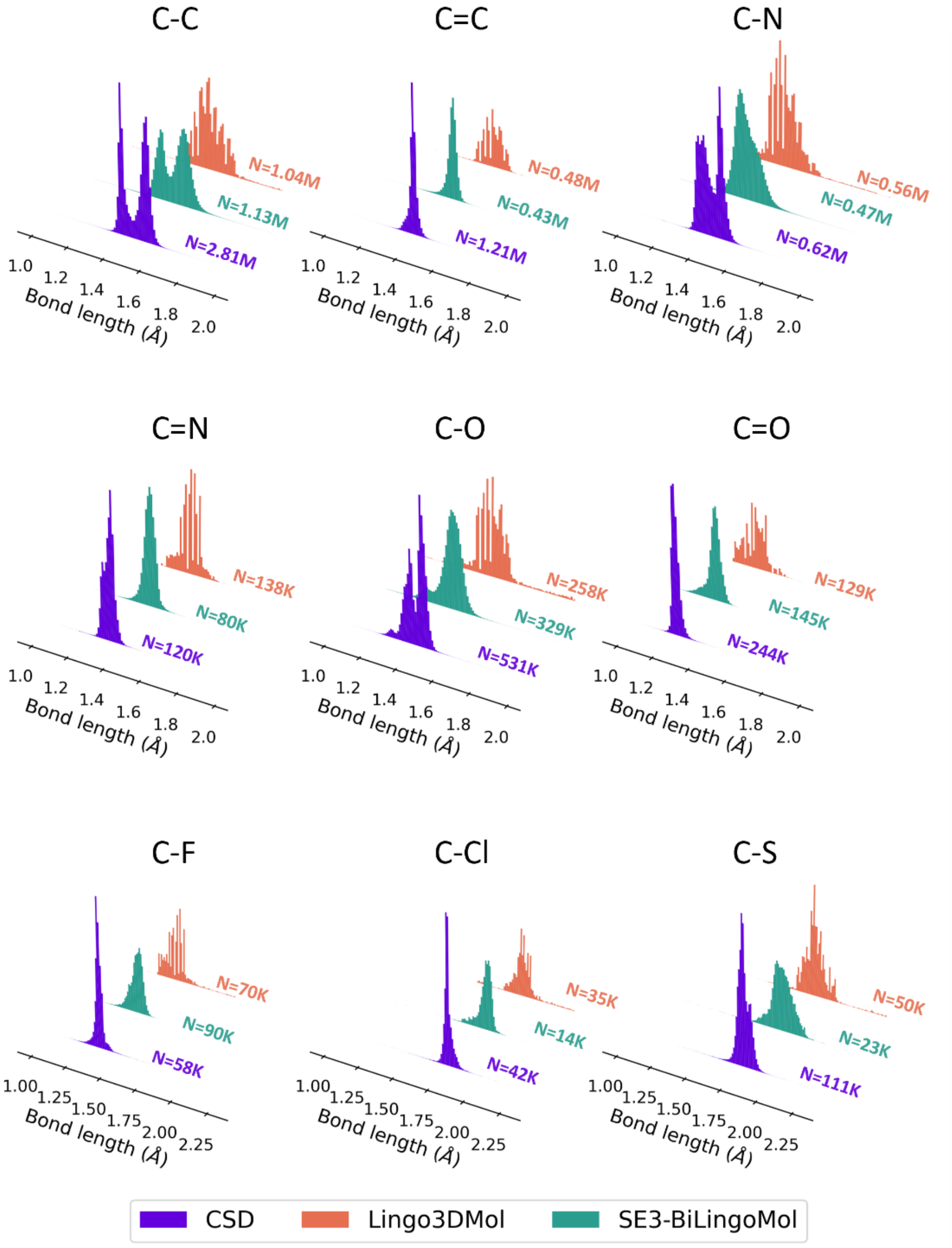

**Extended Data Figure 2. Impact of continuous coordinate representation on bond length distributions.** Histograms illustrating the distributions of nine common bond lengths from experimentally determined structures (CSD, purple), molecules generated by Lingo3DMol (red), and molecules generated by SE3-BiLingoMol (green). Specifically, distributions are shown for C-C, C=C, C-N, C=N, C-O, C=O, C-F, C-Cl, and C-S bonds. Lingo3DMol, which relies on discrete coordinate representation, exhibits jagged and discontinuous bond length distributions. In contrast, our SE3-BiLingoMol model, employing a regression-based continuous coordinate representation, yields smooth and continuous bond length destitutions that closely mirror experimentally determined structures from CSD. The number of molecules analyzed for each specific bond length distribution is indicated by 'N' within the respective histogram.

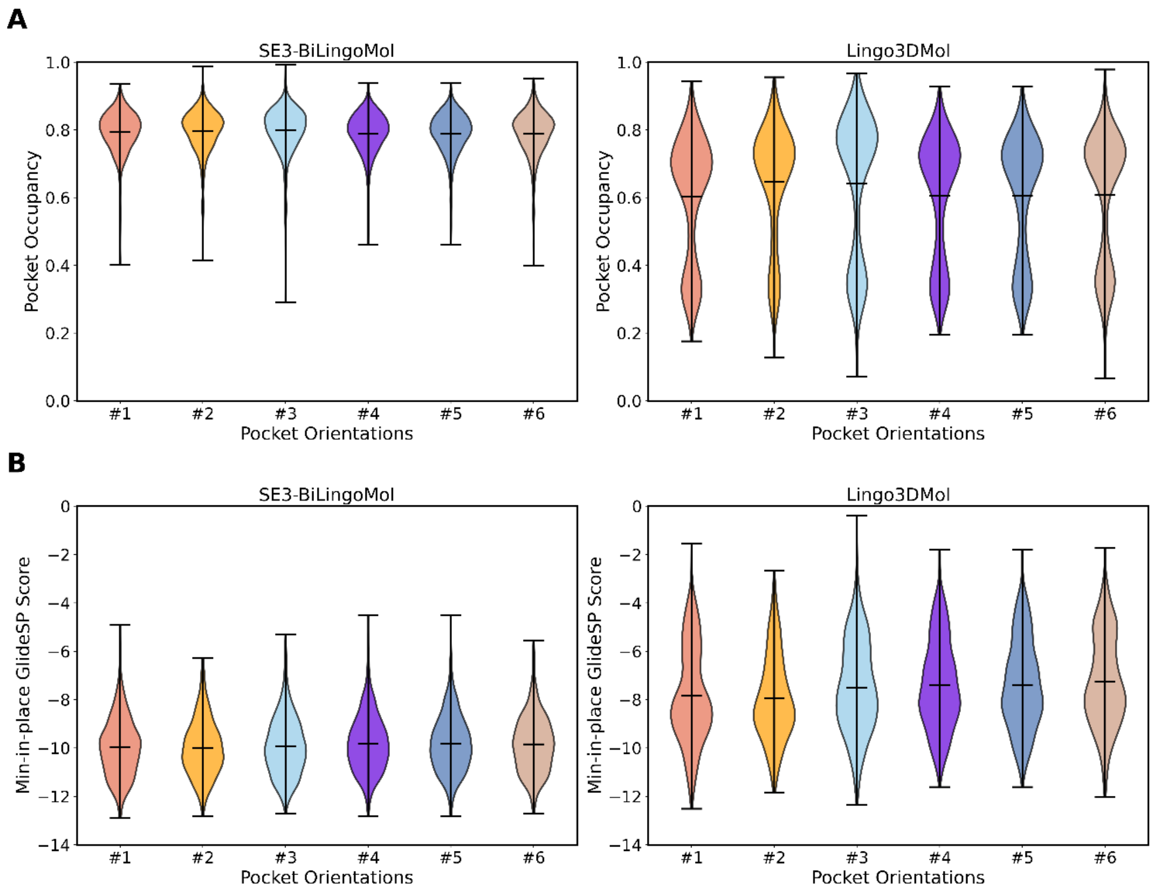

**Extended Data Figure 3. SE(3)-Equivariance Confers Robustness to Symmetry Variations in Molecule Generation: A PDB 2HV5 Case Study.** For this case study, the protein pocket structure from PDB 2HV5 was subjected to six different random rigid-body rotations and translations, creating six distinct pocket orientations. For each of these pocket orientations, 1,000 molecules were generated independently by SE3-BiLingoMol (our SE(3)-equivariant model) and Lingo3DMol (a non-SE(3)-equivariant model).

(A) Violin plots compare the distributions of pocket occupancy for the generated molecules. (B) Violin plots compare the distributions of min-in-place GlideSP scores for the generated molecules.

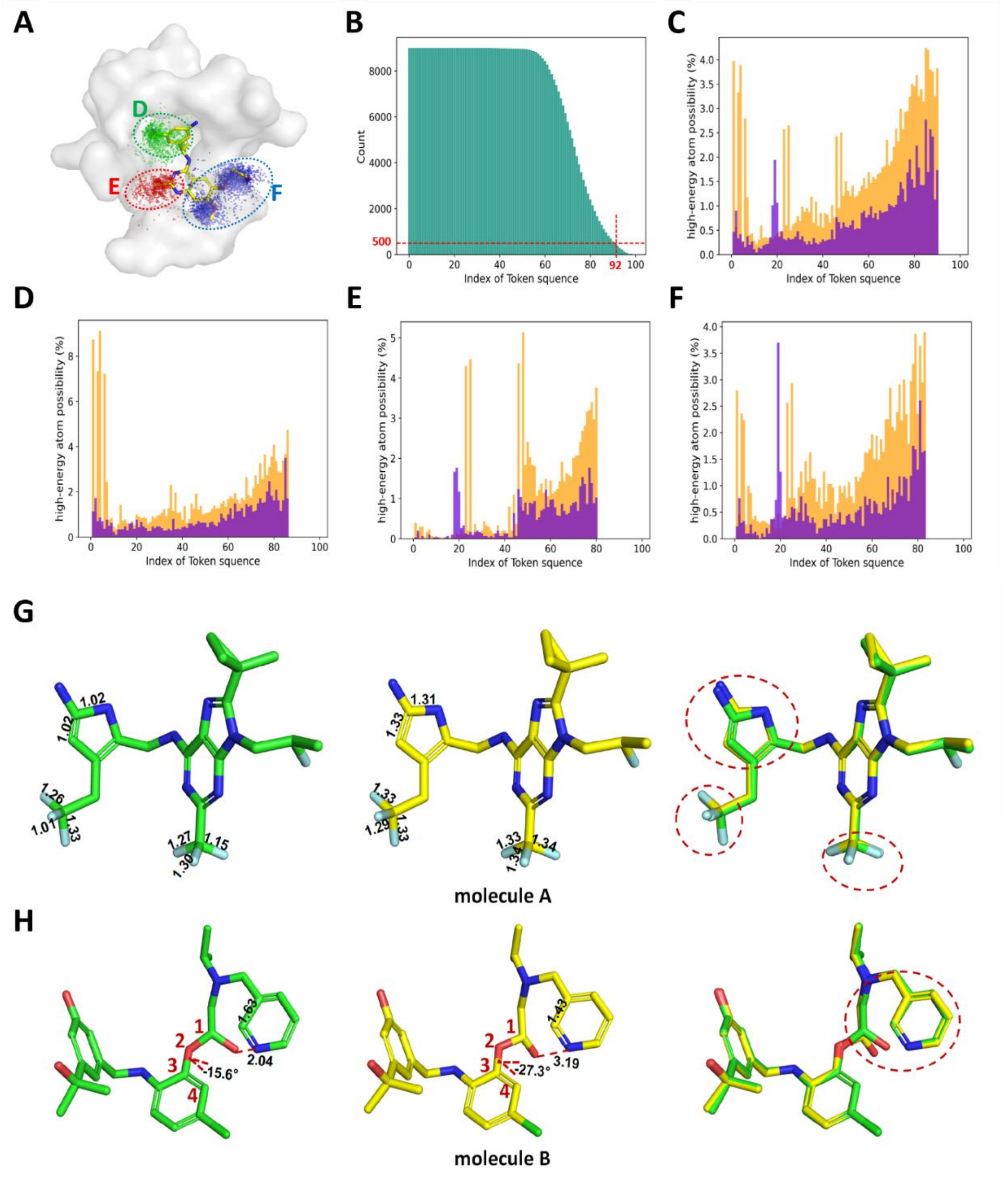

**Extended Data Figure 4. Self-refinement module mitigates conformational errors in autoregressive sampling: A case study**

**(A)** Ligand binding pocket structure from PDB 6SCM. Dots represent the positions of the last sampled atoms from 9,000 generated molecules. These dots are spatially categorized: green indicates that the last atom is deeply located within the binding pocket, while red or blue denote those in two distinct solvent-exposed regions (corresponding to the analyses in panels E and F). Each colored category comprises 3,000 dots.

**(B)** Histogram illustrating the distribution of generated molecules by the number of completed sampling steps. A red dashed line at 500 molecule counts on the vertical axis establishes a minimum sample size for robust observation. This threshold dictates that the high-energy atom (HEAD) analysis in subsequent panels is presented only for sampling steps up to 92, as steps beyond this point involve fewer than 500 molecules that have completed them, thus possibly precluding meaningful interpretation of trends.

**(C)** Bar chart displaying the probability of high-energy atom occurrence (identified by HEAD analysis) for each sampling step. The HEAD method assess conformation quality by quantifying energy anomalies at the atomic level. Data are presented for all generated molecules before (orange) and after (purple) self-refinement, demonstrating a substantial reduction in identified errors.

**(D-F)** Analogous chart depicting conformational error probability before (orange) and after (purple) self-refinement, specifically for molecules whose last atom terminates in distinct regions: (D) deeply inside the pocket; (E) within the red solvent-exposed region (as defined in panel A); or (F) within the blue solvent-exposed region (as defined in panel A). These panels collectively illustrate the consistent amelioration of conformational errors by the self-refinement module across diverse sampling termination environments.

**(G-H)** Representative examples illustrating the correction of initially flawed molecular conformations by the self-refinement module. Initial and refined conformations are depicted in green and yellow, respectively. In panel **(G)**, the presented example initially showed two compressed C-F bonds (1.01 Å and 1.15 Å, compared to a typical 1.35 Å) and a severely distorted pyrrole group. Refinement successfully relaxed the C-F bonds to approximately 1.34 Å and reduced the pyrrole distortion. The example in panel **(H)** initially exhibited a severe steric clash (O-N distance 2.04 Å) and an elongated C-C bond (1.63 Å) within the pyridine ring. Refinement resolved this steric clash by adjusting the C1-O2-C3-C4 dihedral angle and optimized the C-C bond to 1.43 Å.

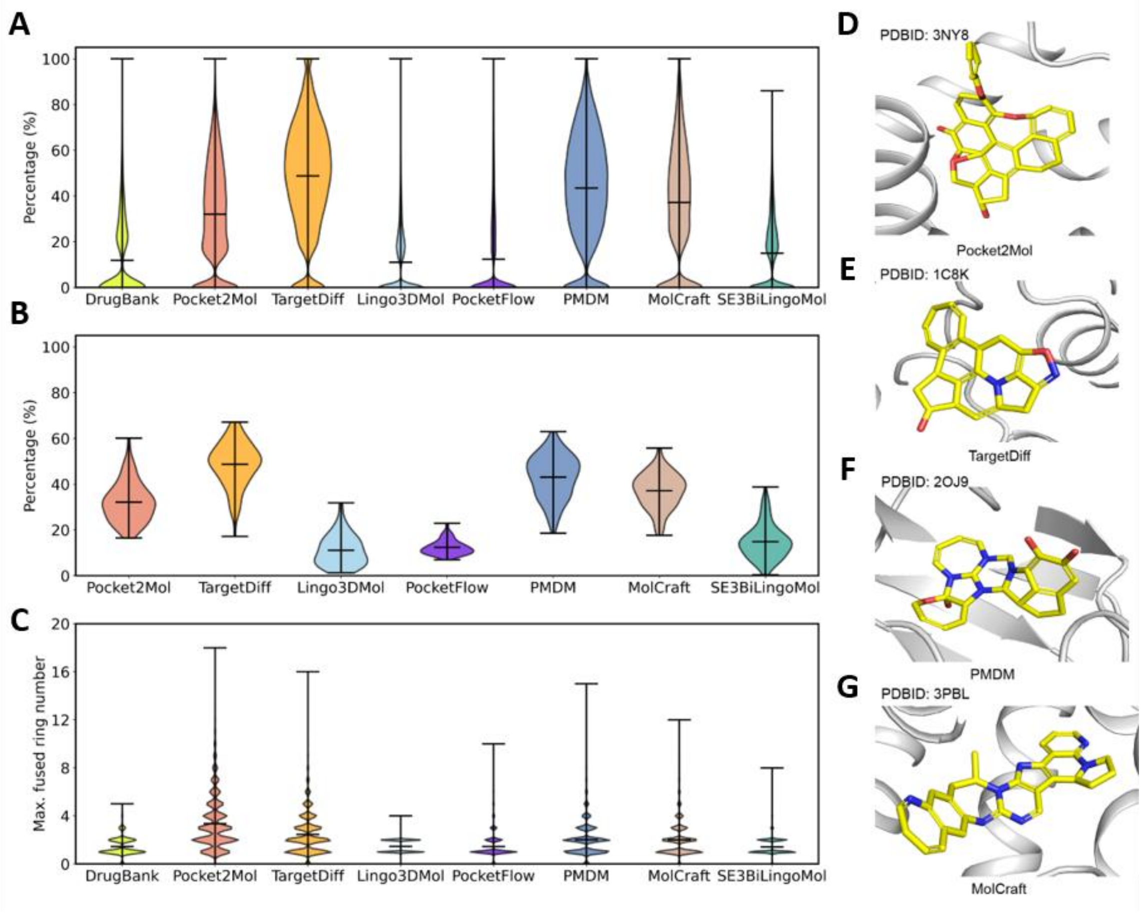

**Extended Data Figure 5. Distributions of non-aromatic ring percentage and maximum fused ring number for the molecules generated by different models.** Drug molecules from DrugBank, filtered with a more stringent drug-likeness criteria (QED > 0.5, SAS < 3, and MW < 700), served as a reference (N=3,743).

**(A)** Distributions for the percentage of non-aromatic rings of all generated molecules.

**(B)** Distributions for the per target average percentage of non-aromatic rings across 102 targets. The DrugBank reference is not applicable and thus excluded, as it lacks explicit target information required for per-target average value calculations.

**(C)** Distributions of maximum number of the fused ring existed in each molecule.

**(D-G)** Cases of generated molecules that contain significant fused ring structures by Pocke2Mol **(D)**, TargetDiff **(E)**, PMDM **(F)** and MolCraft **(G)**.
